# High-yield enzymatic synthesis of mono– and trifluorinated alanine enantiomers

**DOI:** 10.1101/2023.11.28.569005

**Authors:** Manuel Nieto-Dominguez, Aboubakar Sako, Kasper Enemark-Rasmussen, Charlotte Held Gotfredsen, Daniela Rago, Pablo I. Nikel

## Abstract

Fluorinated amino acids are a promising entry point for incorporating new-to-Nature chemistries in biological systems. Hence, novel methods are needed for the selective synthesis of these building blocks. In this study, we focused on the enzymatic synthesis of fluorinated alanine enantiomers. To this end, the alanine dehydrogenase from *Vibrio proteolyticus* and the diaminopimelate dehydrogenase from *Symbiobacterium thermophilum* were applied to the *in vitro* production of (*R*)-3-fluoroalanine and (*S*)-3-fluoroalanine, respectively, using 3-fluoropyruvate as the substrate. Additionally, an alanine racemase from *Streptomyces lavendulae*, originally selected for setting an alternative enzymatic cascade leading to the production of these non-canonical amino acids, had an unprecedented catalytic efficiency in the β-elimination of fluorine from the monosubstituted fluoroalanine. The *in vitro* enzymatic cascade based on the dehydrogenases of *V*. *proteolyticus* and *S*. *thermophilum* included a cofactor recycling system, whereby a formate dehydrogenase from *Pseudomonas* sp. 101 (either native or engineered) coupled formate oxidation to NAD(P)H formation. Under these conditions, the reaction yields for (*R*)-3-fluoroalanine and (*S*)-3-fluoroalanine reached >85% on the fluorinated substrate and proceeded with complete enantiomeric excess. Moreover, the selected dehydrogenases were also able to catalyze the conversion of trifluoropyruvate into trifluorinated alanine, as a first-case example of biocatalysis with amino acids carrying a trifluoromethyl group.

## INTRODUCTION

The extensive array of chemicals essential to our contemporary society is predominantly supplied by the chemical industry. These chemicals include polymers, agrochemicals and pharma molecules, the production of which largely depends on traditional synthetic chemistry. Conventional approaches for the synthesis of these compounds often employ processes and reagents leading to significant environmental hazards, notably through the generation of toxic by-products and waste streams [1,2]. With increasing environmental awareness, there is a growing interest in leveraging Nature and biosynthesis as alternative sources for fulfilling chemical production needs [3,4]. The molecular diversity in the biosphere is considered nearly boundless, yet only a small portion of these biomolecules has been explored so far [5]. Nonetheless, the connection between industrially-relevant chemicals and naturally-occurring products is remarkably limited [6]. A key factor contributing to this disparity is the industrial reliance on synthetic compounds containing chemical elements, e.g. fluorine (F) [7] and other halogen atoms [8], which are not typically found in natural biological systems [9,10]. Establishing a biotechnological alternative to chemical synthesis requires the rational design of biosynthetic pathways and degradation routes either as enzymatic cascades *in vitro* [11,12] or as part of living organisms [13], enabling them to execute new-to-Nature chemistries [14,15].

Amino acids represent an attractive target for bioproduction due to their wide industrial exploitation [16]. The twenty standard proteinogenic amino acids, while relatively simple in structure, are fundamental components of all naturally-occurring polypeptides. Furthermore, the D-enantiomers of some amino acids are essential in the formation of the bacterial peptidoglycan [17] and are involved in the biosynthesis of natural peptide antibiotics [18]. In this sense, introducing the D-enantiomers of non-canonical amino acids (NCAAs) into the chemistry of living bacterial cells represents an effective approach to substantially enhance the chemical diversity of cellular structures [19–21], but the biocatalysis toolbox for their production is somewhat limited [22,23]. Among the broad group of NCAAs, fluorinated amino acids (FAAs), which contain one or more F atoms, have considerable potential for engineering new chemistries. The small size of the F atom renders FAAs structurally similar to their natural analogues, making them largely indistinguishable by the cellular machinery [24]. Currently, most FAAs are produced through chemical synthesis [25], although some of them can be obtained enzymatically *in vitro* [24,26]. Notably, 4-fluoro-L-threonine is a naturally-occurring FAA produced by *Streptomyces cattleya* and other actinomycetes [27]. To date, research approaches for the production of FAAs have primarily focused on the L-enantiomers, exploring their effects when incorporated into individual proteins or at the cellular proteome-level. However, the impact of D-enantiomers of FAAs on the cellular metabolism remains largely unexplored and, to best of our knowledge, there are no reports on the biosynthesis of these compounds at chiral purity.

Alanine (Ala) represents an interesting model for studying the biosynthesis FAAs. The synthesis of Ala occurs through a single enzymatic step involving the reductive amination of pyruvate (Pyr). Owing to the small size of the methyl substituent in this amino acid, the proximity of the F substituent to the amino group allows strong electron-withdrawing effect of the halogen atom to influence the amino group [28], thus making Ala a particularly interesting case study. In terms of structure, glycine is the only amino acid that is simpler than Ala. However, fluorinated derivatives of glycine at the Cα position have been found to be unstable, undergoing rapid defluorination [29]. To date, examples on the bioproduction of 3-fluoroalanine (FAla) have been limited to the L-enantiomer [(*R*)-FAla], which can be obtained through the action of either L-Ala dehydrogenases or ω-transaminases on 3-fluoropyruvate (FPyr) [30–32]. The subsequent formation of the D-enantiomer [(*S*)-FAla] could theoretically be achieved through the action of an alanine racemase (**Scheme 1**). However, this type of enzyme is inhibited by FAla [33], which contributes to the known antibacterial properties of some FAAs. An alternative approach could involve the use of *meso*-diaminopimelate dehydrogenases, enzymes that have garnered interest due to their broad substrate specificity and ability to produce target compounds with high enantiomeric excess [34,35]. *meso*-Diaminopimelate dehydrogenases have proven useful for the biosynthesis of different D-amino acids, including D-Ala [34] and D-phenyalanine [36]. Yet, there are no available reports of their activity on fluorinated derivatives of Pyr or in the production of FAAs—in contrast with the increasing need of synthetic tools to produce these building blocks towards establishing novel chemistries.

In this work, we explored the enzymatic production of mono– and tri-fluorinated Ala from F*_n_*Pyr (with *n* = 1 or 3). To this end, we recombinantly produced an alanine dehydrogenase from *Vibrio proteolyticus* (*^Vp^*ALDH) and a diaminopimelate dehydrogenase from *Symbiobacterium thermophilum* (*^St^*DAPDH). The kinetics of these two enzymes against FPyr and 3,3,3-trifluoropyruvate (F3Pyr) were determined *in vitro*, together with the enantiomer distribution of the corresponding F*_n_*Ala produced. An efficient NAD(P)H regeneration cycle, based on a NAD^+^-dependent formate dehydrogenase (FDH) from *Pseudomonas* sp. 101 and an engineered derivative that exhibits increased specificity for NADP^+^, was implemented to support high-yield bioproduction of FAla. This approach boosted the reaction yields to ∼90% and 18 mM for the D-enantiomer and ∼100% and 20 mM for the L-enantiomer of the FAA. Furthermore, an alanine racemase from *Streptomyces lavendulae* (*^Sl^*AlaR), originally selected as a candidate for assembling these enzymatic cascades, was found to possess an unexpectedly high defluorinating activity on FAla *via* β-elimination. Taken together, the results discussed in this article represent a first-case report on the enzymatic synthesis of both a fluorinated D-amino acid and a trifluorinated version of Ala.

## RESULTS AND DISCUSSION

### Rationale behind enzyme selection, production and purification for establishing *in vitro* synthesis of fluorinated amino acids

In this study, the selection of enzymes was based on their potential to enhance the production of fluorinated alanine (FAla), with an initial round of biocatalyst selection informed by examples in the primary literature. Alanine dehydrogenase, derived from *Vibrio proteolyticus* (*^Vp^*ALDH, a NAD^+^-dependent dehydrogenase), was chosen due to its well-documented ability to maintain >70% substrate specificity for 3-fluoropyruvate (FPyr) in comparison to its non-fluorinated analogue, pyruvate (Pyr) [31]. The diaminopimelate dehydrogenase from *Symbiobacterium thermophilum* (*^St^*DAPDH, a NADP^+^-dependent enzyme), on the other hand, was selected based on its known substrate promiscuity [34]. The Ala racemase of *Streptomyces lavendulae* (*^Sl^*AlaR) was picked owing to its reported resistance to inactivation by D-cycloserine (4-amino-3-isoxazolidinone) [37]—a molecule that, similarly to FAla, can inhibit AlaR by forming an adduct with the pyridoxal 5′-phosphate (PLP) cofactor and a key catalytic lysine residue in the active site [38]. The particular architecture of the active site of *^Sl^*AlaR led to the hypothesis that this enzyme variant should be resistant to D-cycloserine inhibition, hence, it was hypothesized that this AlaR variant might also exhibit increased tolerance towards FAla, an essential requirement in this study (**Scheme 1**). Additionally, an Ala racemase from *Escherichia coli* (*^Ec^*AlaR) was adopted as a reference, since its inhibition by FAla has been previously characterized [39]. Finally, a NAD^+^-dependent formate dehydrogenase (FDH) from *Pseudomonas* sp. 101 (NAD-*^Pse^*FDH) and an engineered variant with increased NADP^+^ preference [40] were incorporated in the designs towards efficient regeneration of NAD(P)H, cofactors required by both *^Vp^*ALDH and *^St^*DAPDH. All the enzymes were successfully produced in *E. coli* BL21(DE3) from the corresponding synthetic DNA fragments and purified to homogeneity through one-step Ni^2+^-nitriloacetic (NTA) affinity chromatography. SDS-PAGE analysis (**Supplementary Fig. S1**) showed that all the proteins were isolated with a high degree of purity.

### Synthesis of FAla and F_3_Ala from the corresponding pyruvate substrates by two dehydrogenases from *Vibrio proteolyticus* and *Symbiobacterium thermophilum*

The initial observation that alanine dehydrogenases (ALDH) are capable of producing (*R*)-FAla through the reductive amination of FPyr was documented in the seminal work of Ohshima et al. [30]. Notwithstanding, this study predominantly focused on the affinity of this enzyme for halogenated substrates, yet a comprehensive kinetic characterization of *^Vp^*ALDH was missing not only in the original report but also in subsequent studies employing this enzyme as a biocatalyst [31,41,32]. Similarly, the broad substrate specificity of *^St^*DAPDH has been underscored in the report by Gao et al. [34], yet experimental assays specifically targeting FPyr have not been conducted for this enzyme or for any other biocatalyst within the *meso*-diaminopimelate dehydrogenase family. Furthermore, there is a dearth of studies exploring the enzymatic conversion of F3Pyr into the corresponding trifluorinated Ala—a new-to-Nature building block with potential for a range of biocatalysis applications [42]. To bridge this knowledge gap, particularly regarding the production of FAla *via* reductive amination, and to evaluate the potential application of this process *in vivo*, an in-depth analysis of the kinetics of both enzymes was executed *in vitro*. The experimental design involved using Pyr, FPyr and F_3_Pyr as substrates. The enzymatic activities were quantified through continuous spectrophotometric monitoring of NAD(P)H oxidation according to the enzymatic reaction in the reductive substrate amination direction, as indicated for pathways involving either ALDH (**Scheme 1**) or DAPDH (**Scheme 2**). The kinetic data were analyzed through the canonical Michaelis-Menten model (**Fig. 1A**).

**Fig. 1.**
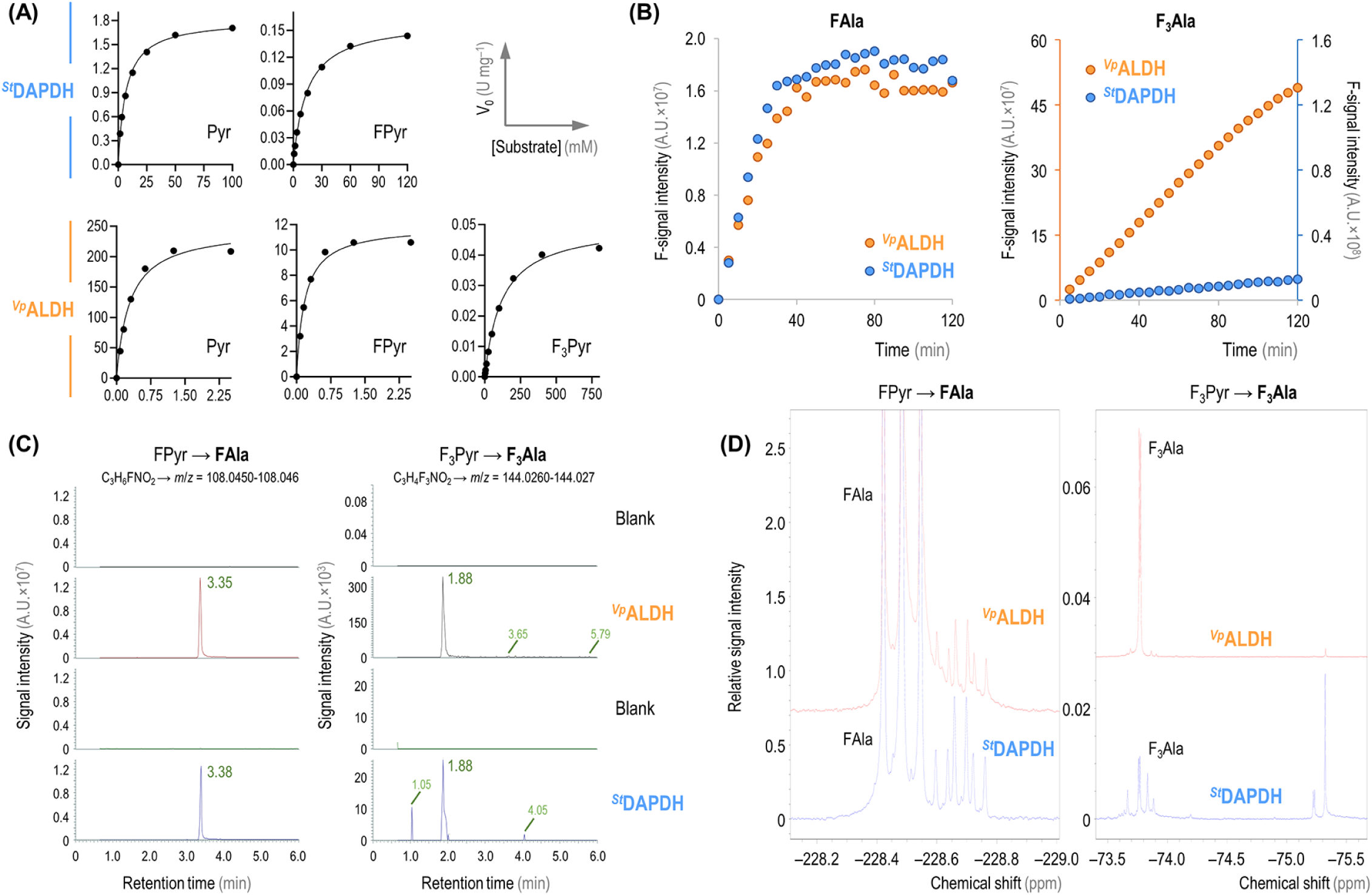
Evaluating the activity of *^St^*DAPDH and *^Vp^*ALDH *in vitro*. **(A)** Kinetic data for the two enzymes assayed against Pyr, FPyr and F_3_Pyr, with fitting to the canonical Michaelis-Menten equation. **(B)** Time-resolved ^19^F-NMR monitoring of FAla and F_3_Ala production. The plots show the integration of the product peak over time for both *^Vp^*ALDH and *^St^*DAPDH; due to the disparity on the levels of F_3_Ala formation with the later enzyme, a secondary *y*-axis is shown (identified with the same color code as per the enzyme assayed). *A.U.*, arbitrary units. **(C)** LC-MS spectra indicating the peaks with the *m/z* signal predicted for FAla (*left*) and F_3_Ala (*right*). The results shown for *^Vp^*ALDH and *^St^*DAPDH are compared to the corresponding blank assay with no added enzyme. *A.U.*, arbitrary units. **(D)** Representative ^19^F-NMR spectra for the assays producing FAla (*left*) and F_3_Ala (*right*) using the corresponding enzymes and substrates. The plots display a zoom-in of the chemical shifts predicted for each of the fluorinated products. The spectrum for F_3_Ala formation by *^St^*DAPDH was acquired using 4 times more scans than for *^Vp^*ALDH in order to increase the signal intensity.

*^Vp^*ALDH showed a similar affinity for the non-fluorinated and the monofluorinated Pyr substrate (*Km* = 0.29 and 0.18 mM, respectively; **Table 1**), but the presence of F mediated a dramatic decrease in the dehydrogenase activity (i.e. only ca. 5% of the *V_max_* was retained when using FPyr as substrate). This result seems to contradict the >70% specific activity on FPyr previously reported for this enzyme [31], probably attributable to differences in the assay setup (e.g. substrate concentration). The *V_max_* value attained in our present study, however, is close to the ∼6% reported for the ALDH enzyme from *Helicobacter aurati* when acting on fluorinated Pyr [32]. One way or the other, *k_cat_*/*Km* values were reduced by >10-fold (from ca. 597 to 46 s^−1^ mM^−1^) when FPyr was used as the substrate. F_3_Pyr, on the other hand, had a strong detrimental effect on both the substrate affinity (*Km* = 121 mM) and the activity (ca. 0.05 U mg^−1^) of *^Vp^*ALDH (**Table 1**). The strong negative impact on the enzyme activity might be due to the polarizing effect of the three F atoms (in the CF_3_ group), which are expected to substantially modify the formal distribution of charges in their molecular surroundings [43]. Conversely, the loss of substrate affinity might seem more surprising, since the small size of F is not predicted to alter the intramolecular structure significantly or cause steric hindrances at the catalytic pocket [9]. However, the analysis of the crystal structure of the ALDH enzyme of *Mycobacterium tuberculosis* indicates that the stabilization of Pyr in the active site depends on the formation of specific hydrogen bonds and charge interactions through three conserved amino acid residues [44], a conformational architecture that might be disturbed by the polarizing effect of the three F substituents in F_3_Pyr. The catalytic residues involved in these interactions (Arg15, Lys75 and His96) were confirmed to play the same stabilizing role in the *^Gk3448^*ALDH enzyme of *Geobacillus kaustophilus* [45], and they are all conserved in *^Vp^*ALDH (**Supplementary Fig. S2**). Prompted by these results, we also analyzed the catalytic performance of *^St^*DAPDH under the same reaction conditions (**Fig. 1A**). The affinity of *^St^*DAPDH for the non-fluorinated, native substrate was ca. 23-fold lower than that of *^Vp^*ALDH; the *k_cat_* values were similarly lower for the dehydrogenase of *S*. *thermophilum*. In general, the trend observed in the *Km* and *V_max_* values evaluated in the presence of fluorinated substrates was similar to the results obtained for the *^Vp^*ALDH enzyme. The effect of FPyr was especially noticeable at the activity level, while the affinity was kept within the same order of magnitude. The *k_cat_*/*Km* was reduced by ca. 1,000-fold in the presence of FPyr (ca. 7×10^−3^ s^−1^ mM^−1^). Moreover, in this case, the reduction of both affinity and activity when using F_3_Pyr as the substrate was too strong to allow for the determination of enzyme kinetic parameters (**Table 1**).

**Table 1.** *In vitro* kinetic characterization of *^Vp^*ALDH and *^St^*DAPDH.

| Enzyme | Substrate | Kinetic parameter |  |  |  |
| --- | --- | --- | --- | --- | --- |
| | | $K_m$ (mM) | $V_{max}$ (U mg<sup>-1</sup>) | $k_{cat}$ (s<sup>-1</sup>) | $k_{cat}/K_m$ (s<sup>-1</sup> mM<sup>-1</sup>) |
| $V_p$ALDH | Pyr | 0.29 ± 0.04 | 247 ± 11 | 173 ± 8 | 596.6 |
|  | FPyr | 0.18 ± 0.02 | 11.9 ± 0.4 | 8.3 ± 0.3 | 46.1 |
|  | F<sub>3</sub>Pyr | 121 ± 8 | 0.051 ± 0.001 | 3.53×10<sup>-2</sup> ± 8×10<sup>-4</sup> | 2.9×10<sup>-4</sup> |
| $S^t$DAPDH | Pyr | 6.7 ± 0.3 | 1.81 ± 0.02 | 1.05 ± 0.01 | 0.16 |
|  | FPyr | 14.0 ± 0.6 | 0.161 ± 0.002 | 0.093 ± 0.001 | 6.6×10<sup>-3</sup> |
|  | F<sub>3</sub>Pyr | N.D. | N.D. | N.D. | N.D. |

The parameters reported in the table correspond to regression coefficients obtained by fitting the mean values from three independent experiments to the Michaelis-Menten model ± standard deviation of the regression coefficient estimates. *N.D.*, not detected.

To facilitate a direct and quantitative assessment of the formation of the target fluorinated products, serving as a supplementary method to the *in vitro* assays (**Fig. 1A**), individual reactions were prepared under the same conditions described above and analyzed by both ^19^F-NMR (**Fig. 1B** and **1D**) and high-resolution LC-MS(MS) (**Fig. 1C**). In the case of the ^19^F-NMR assays, the dynamics of both substrate depletion and product generation in the reactions were continuously monitored in real-time. This combined analytical approach provided a deeper understanding of the reaction kinetics and enabled the direct visualization of the transformation of the substrates within the reaction milieu. Both FPyr and F_3_Pyr could be easily identified by ^19^F-NMR (**Supplementary Fig. S3A** and **S3B**, respectively). In general, the chemical shifts and the multiplet patterns observed in these samples matched the expected signals for both FAla and F_3_Ala, regardless of the selected enzyme (*^Vp^*ALDH or *^St^*DAPDH, **Fig. 1D**). FAla formation, evaluated as the time-resolved intensity of the F-signal in the samples, reached a plateau around 40 min of incubation for both enzymes (**Fig. 1B**). Conversely, the transformation of F_3_Pyr followed a linear kinetic over 2 h, with a significantly lower conversion rate mediated by the action of *^St^*DAPDH (**Fig. 1B**), suggesting that the lower rates observed for this substrate prevented the reactions to reach equilibrium within the duration of the assay. These results recapitulate the experimental observations in the assays where product formation was inferred from the rates of NADH oxidation (**Table 1**).

To further verify the chemical identity of the expected FAAs, the samples were also submitted to LC-MS(MS) analysis in positive-heated electrospray ionization (HESI) mode (**Fig. 1C**). While no significant signal could be detected in the blank (control) experiments, the reaction samples displayed signals with an experimental *m*/*z* fitting the predicted values for FAla and F_3_Ala—for FAla (C_3_H_6_FNO_2_), *m*/*z* = 108.045-108.046 and for F_3_Ala (C_3_H_6_F_3_NO_2_), *m*/*z* = 144.026-144.027 (**Fig. 1C**). The predicted masses for both enzymes and substrates were likewise confirmed in these assays. An MS(MS) validation was successfully carried out for every condition except for F_3_Ala production by *^St^*DAPDH, due to limited signal intensity. The full MS(MS) spectra for the conversion of FPyr into FAla by both *^Vp^*ALDH and *^St^*DAPDH are presented in **Supplementary Fig. S4A** and **S4B**, respectively; while the MS(MS) spectrum for the transformation of F_3_Pyr by *^Vp^*ALDH is shown in **Supplementary Fig. S5**. The ^19^F-NMR spectra further confirmed the identity of the target fluorinated products, with relative intensities considerably higher for FAla than F_3_Ala (**Fig. 1D**). Accordingly, the spectra for the trifluorinated product were acquired using 4 times more scans in order to increase the signal intensity. Interestingly, the analysis of the performance of *^St^*DAPDH against F_3_Pyr as a substrate revealed the formation of side products with signal intensities within the range of (or even higher than) that expected for F_3_Ala (**Fig. 1D**). Both the chemical shift and the singlet nature of these signals are suggestive of the presence of a –CF_3_ group. Since these fluorinated side products are virtually absent in the equivalent reaction mixture with *^Vp^*ALDH as the biocatalyst, they are hypothesized to be the result of an alternative catalytic fate in the active site of *^St^*DAPDH [46,47].

### An alanine racemase from *Streptomyces lavendulae* displays a secondary dehalogenation activity on fluorinated amino acid analogues

As indicated in the previous section, *^Vp^*ALDH had the best catalytic efficiency for the reductive amination of FPyr, with a *k_cat_*/*Km* ca. 7,000-fold higher than *^St^*DAPDH (**Table 1**). However, (*R*)-FAla is the predicted product for the *^Vp^*ALDH activity, which requires the action of a racemase to convert the enantiomer product into (*S*)-FAla. Hence, an experimental screening was established to investigate if *^Sl^*AlaR is able to both escape the inhibition mediated by FAla while catalyzing the formation of the desired enantiomer. As a first step, the *^Sl^*AlaR and *^Ec^*AlaR racemases were assayed against its natural substrate, L-Ala, by coupling the reaction to the Pyr-dependent formation of L-lactate (**Supplementary Scheme S1**). The reaction was monitored by following the rate of NAD^+^ generation, which indicated that both *^Sl^*AlaR and *^Ec^*AlaR were active under these conditions (**Supplementary Fig. S6A** and **S6B**, respectively). Next, a similar spectrophotometric assay was designed to couple the activity of AlaR with the oxidation of NADH in a different configuration (**Fig. 2A**). In this case, the activity of *^Vp^*ALDH on FPyr was combined with *^Sl^*AlaR and a D-amino acid oxidase (DAAO, a commercially-available DAAO from porcine kidney). DAAO displays specific activity on the D-enantiomer, and this enzyme was expected to regenerate FPyr while producing H_2_O_2_ as byproduct of the deamination reaction. Owing to the formation of this byproduct, which could potentially inhibit the enzymes if the peroxide concentration increases above a threshold level, a catalase from bovine liver was added to the reaction mixture. In these tests, NADH was added in excess (1 mM) in comparison to the FPyr concentration (0.5 mM) as a strategy to study the functionality of *^Sl^*AlaR through the formation of NAD^+^ while preventing any cofactor limitation. The underlying reasoning was that the reaction could only proceed to completion (i.e. NADH exhaustion) only if the DAAO activity effectively replenished FPyr (**Scheme 3**). Control experiments were prepared with either no enzymes added, *^Vp^*ALDH alone and also combining *^Vp^*ALDH and *^Sl^*AlaR but without DAAO. The latter assay was set to account for the potential β-elimination activity of the AlaR enzyme, which would lead to the release of non-fluorinated Pyr, thus creating an alternative regeneration cycle (**Fig. 2A**). Equivalent reaction mixtures, using an excess of FPyr (5 mM), were prepared as additional controls (**Supplementary Fig. S7**). In these reactions, all NADH seemed to be consumed at similar rates in < 40 min across the experimental conditions tested.

**Fig. 2.**
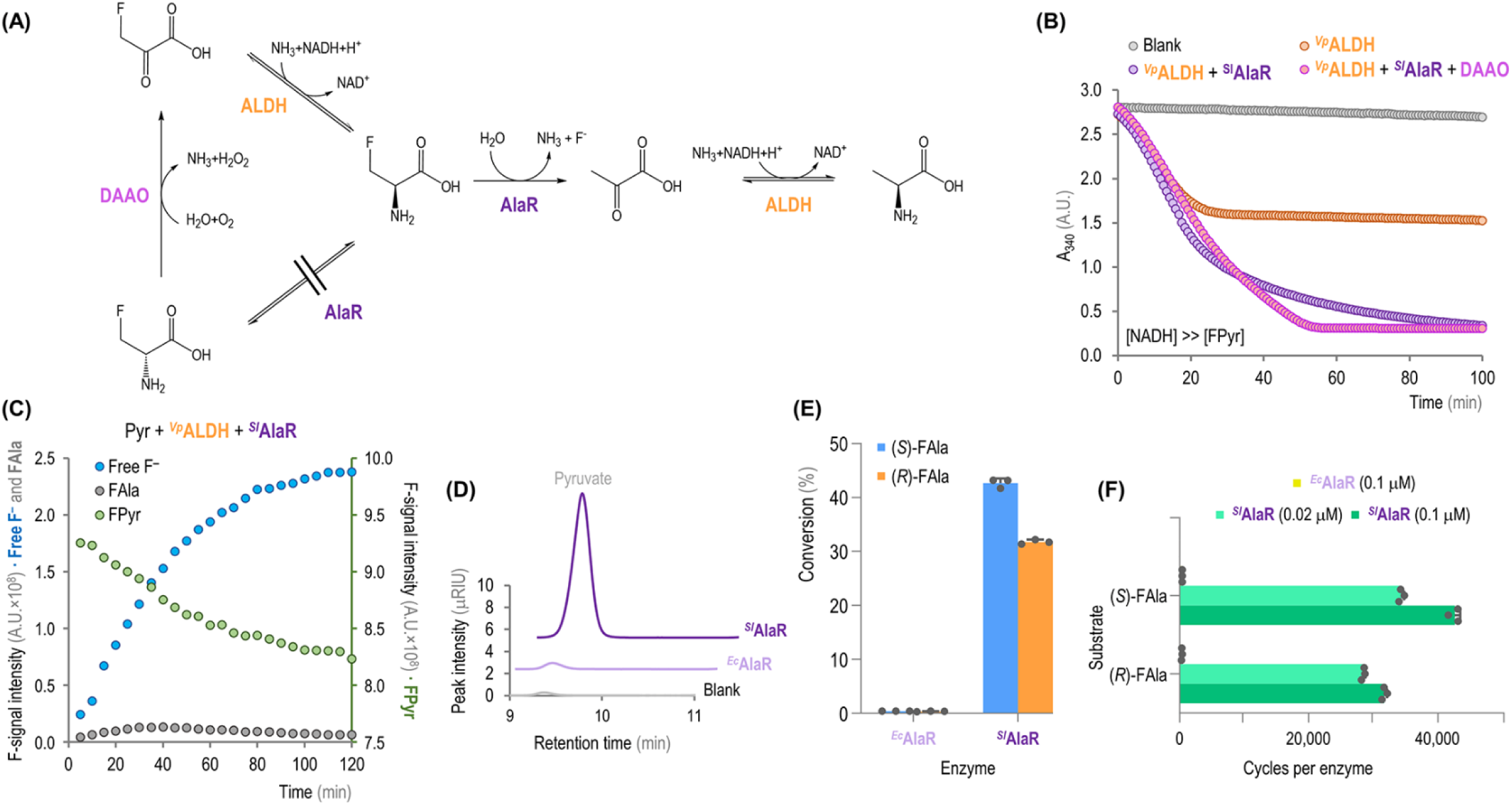
Exploring the dehalogenation activity of *^Sl^*AlaR. **(A)** The scheme depicts the possible pathways for the synthesis of an ALDH substrate, which can be formed through β-elimination of the F atom or *via* the action of a racemase upon the addition of DAAO. **(B)** Spectrophotometric monitoring of NADH oxidation using FPyr as limiting substrate with different enzyme combinations. Each point represents the mean value from three independent experiments. *A.U.*, arbitrary units. **(C)** Time-resolved ^19^F-NMR monitoring of FAla formation, FPyr consumption and release of free fluoride (F^−^). The axes are colored according to the chemical species measured in each case. *A.U.*, arbitrary units. **(D)** Representative example of the HPLC profiles for assays incubated with either *^Sl^*AlaR or *^Ec^*AlaR and the blank for the same reaction conditions. The plot shows the region encompassing the characteristic retention time of Pyr. *μRIU*, refractive index units × 10^− 6^. **(E)** Substrate conversion per enantiomer [(*S*)– and (*R*)-FAla], determined from the release of Pyr as compared to the initial amount of FAla in the assays. **(F)** Catalytic cycles per unit of enzyme for both substrates and different enzyme amounts. In panels **(E)** and **(F)**, the bar graphs represent mean values ± standard deviations from three independent experiments, individual data points are indicated in the plots.

Surprisingly, in the reactions where FPyr was the limiting substrate, the combination of *^Vp^*ALDH and *^Sl^*AlaR was sufficient to reach similar NADH oxidation levels as those observed in assays containing *^Vp^*ALDH, *^Sl^*AlaR and DAOO (**Fig. 2B**). As expected, NAD^+^ formation was rather limited in assays containing *^Vp^*ALDH alone (ca. half of the NADH was oxidized over the first 30 min of the test, indicative of the conversion of FPyr into FAla without any further transformation taking place). These results suggested that the β-elimination activity of *^Sl^*AlaR had to be much higher than the levels reported for its orthologue from *E. coli*, an AlaR enzyme thoroughly characterized *in vitro* [39,48]. To further substantiate this hypothesis, the dynamics of the fluorinated substrate (FPyr) and potential product(s) thereof were monitored by ^19^F-NMR in assays combining *^Vp^*ALDH and *^Sl^*AlaR (**Fig. 2C** and **Supplementary Fig. S8**). This time-resolved experiment revealed a steady release of free fluoride (F^−^) from the substrate that essentially mirrored the decrease in the FPyr-related F-signal, while the amount of FAla remained at background levels throughout the 2 h assay.

To explore the extent of the defluorination capacities of *^Sl^*AlaR, a direct assay was prepared by testing the activity of this enzyme against the pure enantiomers, i.e. (*R*)-FAla and (*S*)-FAla. Equivalent reactions were prepared with *^Ec^*AlaR to be used as a reference, on the basis of the well-characterized inhibition of this enzyme in the presence of either FAla enantiomer. In these assays, the β-elimination activity of the AlaR racemases was evaluated through the quantification of the released Pyr by HPLC, which is stoichiometric with respect to the formed F^−^ *via* β-elimination. The HPLC profile of these reactions indicated that Pyr was indeed formed in the reactions containing *^Sl^*AlaR, while residual amounts of the non-fluorinated product could be detected in the assays with *^Ec^*AlaR (**Fig. 2D**). Furthermore, (*R*)-FAla and (*S*)-FAla were transformed by *^Sl^*AlaR into Pyr at ca. 32% and 43% conversion yields, respectively (**Fig. 2E**). Conversely, *^Ec^*AlaR had a very low level of β-elimination activity (paired to the inhibitory effect of fluorinated substrates), and only <0.5% substrate conversion was detected for both FAAs.

The number of catalytic cycles per enzyme unit (Ψ) was also explored for the two racemase enzymes under study (**Fig. 2F**). Two biocatalyst concentrations (0.02 and 0.1 μM) were used in these assays in order to cover a 5-fold range of enzyme/substrate ratios, and the Ψ values were determined as molecules of product per molecule of enzyme. Under these assay conditions, Pyr formation was below the experimental limit of detection when *^Ec^*AlaR was used at 0.02 μM. When the biocatalyst concentration was increased to 0.1 μM, *^Ec^*AlaR mediated Ψ = 400 cycles for (*S*)-FAla and Ψ = 300 cycles for (*R*)-Fala (**Fig. 2F**). *^Sl^*AlaR, on the other hand, reached Ψ = 34,000-42,000 cycles for (*S*)-FAla and Ψ = 28,000-31,000 cycles for (*R*)-FAla when the racemase was assayed at 0.02 and 0.1 μM, respectively (**Fig. 2F**). Therefore, *^Sl^*AlaR had an overall 100-fold increased capacity of mediating FPyr defluorination as compared to *^Ec^*AlaR. In this sense, the number of β-elimination cycles prior irreversible inactivation by FAla (both enantiomers) has been experimentally determined for *^Ec^*AlaR [39] and other microbial racemases from *Salmonella* [38,49] and *Pseudomonas* species [50]. In all these documented cases, Ψ = 800 cycles (or even lower), which is substantially lower than our experimental results for *^Sl^*AlaR on this FAA. Although the *in vitro* assay adopted throughout this study was merely designed to measure the number of catalytic cycles per unit of enzyme under selected experimental conditions, our results would indicate that *^Sl^*AlaR exhibits an unprecedent defluorination capacity. This observation may have a functional connection to the fact that *Streptomyces* species are known to host reactions for fluorometabolite biosynthesis [51], which require exquisite biochemical [52] and physiological adaptations [53] to avoid toxicity. Based on the results described in this section, *^Sl^*AlaR was deemed unsuitable to racemize (*R*)-FAla as required to produce D-FAAs—but it might be repurposed as a biodehalogenation tool in future research.

### *In vitro* NAD(P)H regeneration supports continuous production of FAla enantiomers with high yields and purity

The main technical limitation for the enzymatic production of FAla from FPyr is the high cost of the redox cofactor. Hence, cofactor regeneration strategies were contemplated in an attempt to improve the yield of the target FAA in a cost-effective fashion. In this sense, energy– and redox-cofactor regeneration cycles have been successfully included as part of enzymatic cascades *in vitro* to boost catalytic efficiency and product formation [54–56]. One of the first approaches implemented to solve this stoichiometric bottleneck for FAA synthesis was proposed by Ohshima et al. [30], and it consisted of coupling NADH regeneration to formate oxidation by means of a FDH enzyme from *Saccharomyces cerevisiae*. Not only is formate a low-cost, readily available additive, but the product of the auxiliary reaction is CO_2_, which is spontaneously eliminated from the reaction mixture by evaporation. While this setup works reasonably well with NADH, NADP^+^ reduction by FDH is notoriously difficult. Previous attempts to modify the cofactor specificity of these dehydrogenases came at the cost of a dramatic loss of affinity on the carbon substrate (*Km* = 1,000 mM for formate) [57]. In this work, the FDH from *Pseudomonas* sp. 101 was selected to support redox cofactor cycling (**Scheme 4**). Importantly, an engineered version of this enzyme can efficiently use NADP^+^ as the cofactor while retaining ∼15% of the catalytic efficiency of the wild-type dehydrogenase against formate [40]. Thus, NAD-*^Pse^*FDH (i.e. the original enzyme from *Pseudomonas* sp. 101) was used to regenerate NADH to support the synthesis of (*R*)-FAla from FPyr, catalyzed by *^Vp^*ADLH (**Fig. 3A**). Similarly, NADP-*^Pse^*FDH was adopted to regenerate NADPH for the production of (*S*)-FAla, catalyzed by *^St^*DADPH (**Fig. 3A**). A separate set of *in vitro* assays confirmed the high specificity for either cofactor of the purified NAD-*^Pse^*FDH and NADP-*^Pse^*FDH enzymes (**Supplementary Fig. S9A** and **S9B**, respectively). Both NH_4_Cl and formate were added to these cascade reactions at high concentration (200 mM) to shift the equilibrium towards the reductive amination. The corresponding reactions were analyzed in terms of total product formation and enantiomeric distribution. In this case, a derivatization step with the commercially-available Marfey’s reagent (1-fluoro-2,4-dinitrophenyl-5-L-alanine amide, FDAA) was incorporated into the experimental design to enable the separation of the two FAla enantiomers by reversed-phase chromatography (**Fig. 3B**). Calibration curves, prepared with the authentic (*R*)– and (*S*)-FAla standards treated with the Marfey’s reagent (**Supplementary Fig. S10A** and **S10B**, respectively), were run in parallel for each set of experiments.

**Fig. 3.**
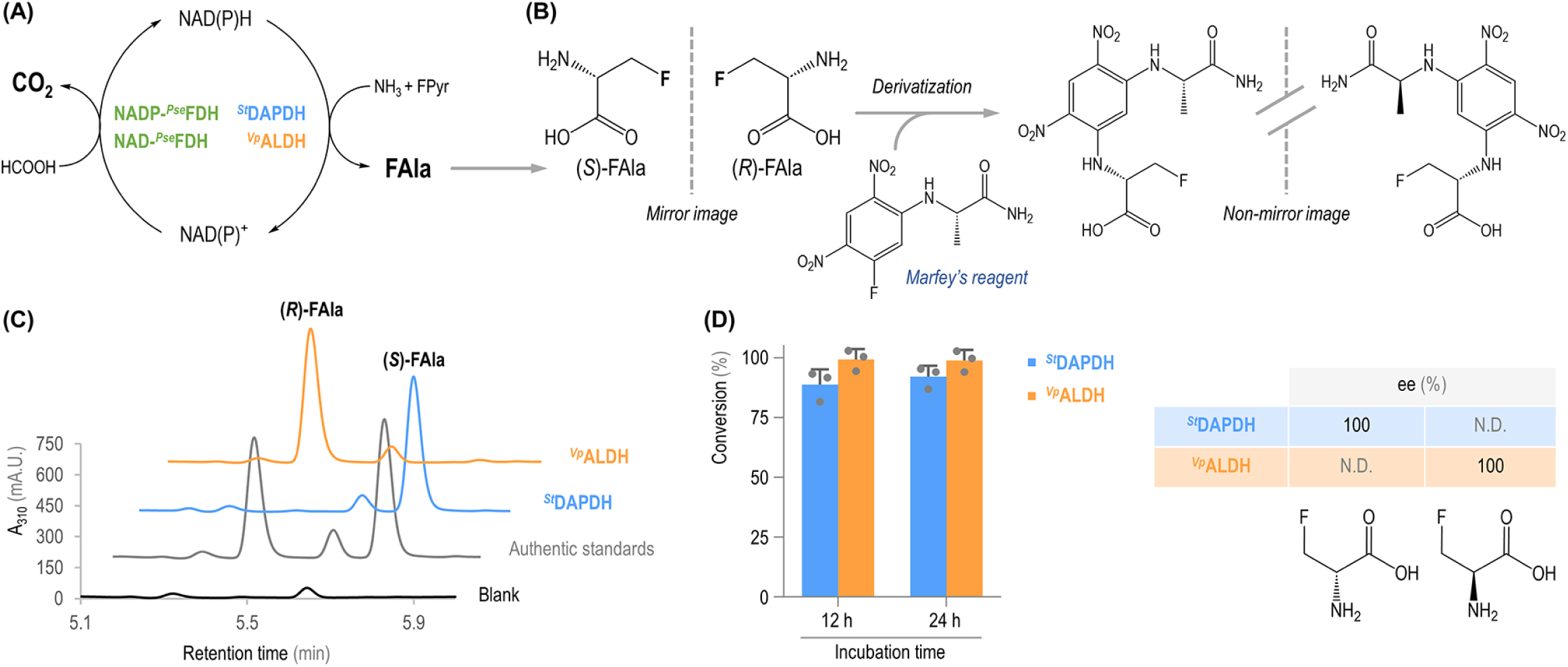
Enzymatic production of FAla *in vitro* and a workflow to determine the enantiomeric excess. **(A)** The scheme indicates the enzymatic cascade assembled for FAla synthesis coupled with the regeneration of the reduced cofactor [NAD(P)H]. The enzyme pair *^St^*DAPDH and NADP-*^Pse^*FDH uses FPyr and formate as substrates; FPyr is converted into (*S*)-FAla by *^St^*DAPDH, whereas NADPH is regenerated by formate oxidation to CO_2_. The equivalent reactions are catalyzed by the enzyme pair *^Vp^*ALDH and NAD-*^Pse^*FDH, in this case forming (*R*)-FAla and providing NADH. **(B)** Chiral purity is assessed after the reaction mixture is derivatized with the Marfey’s reagent (1-fluoro-2,4-dinitrophenyl-5-L-alanine amide, FDAA), which converts the FAla enantiomers into non-enantiomeric derivatives that can be separated by reversed-phase chromatography. **(C)** Representative HPLC profiles showing a reaction blank, a mixture of authentic (*S*)-FAla and (*R*)-FAla standards, a sample of the reaction containing *^St^*DAPDH and a sample with *^Vp^*ALDH. *mA.U.*, arbitrary units × 10^−3^. **(D)** Quantification of the FAla formation in terms of FPyr conversion and enantiomeric excess, ee, for *^Vp^*ALDH and *^St^*DAPDH. The bar graph represents mean values ± standard deviations from three independent experiments, individual data points are indicated in the plot. *N.D.*, not detected.

Analysis of the HPLC profiles indicated that this methodology allowed for the determination of chiral purity (**Fig. 3C**). Moreover, both *^Vp^*ALDH and *^St^*DAPDH had a specific pattern of product formation, mediating the selective synthesis of (*R*)-FAla and (*S*)-FAla, respectively. High-conversion yields were detected for both enzymes, with ∼100% and ∼90% conversion of FPyr into the target FAA products by *^Vp^*ALDH and *^St^*DAPDH, respectively (**Fig. 3D**). These product yield values were similarly high when comparing reactions incubated for either 12 or 24 h, suggesting that the excess of NH_4_Cl and formate was sufficient to keep the reaction equilibrium shifted in the reductive amination direction. Moreover, the enantiomeric excess was 100% for both enzymes (**Fig. 3D**), with no significant detection of the alternative enantiomer form of the FAA— underscoring the very selective nature of the reaction catalyzed by either dehydrogenase. Under these conditions and upon an incubation of 24 h, (*R*)-FAla was produced at 20 mM in the enzymatic cascade containing *^Vp^*ALDH, whereas the concentration of (*S*)-FAla reached 18 mM when *^St^*DADPH was used as the biocatalyst. These values highlight the functionality of the FDH-based regeneration system, without which the maximal production levels would be constrained by the amount of reduced cofactor added (1 mM). Furthermore, the FAla yields on substrate obtained in this study rank among the highest reported for FAAs, while the specific formation of the D-enantiomer had not been reported thus far. While these results were somewhat within the range expected for *^Vp^*ADLH [30], the high performance of *^St^*DADPH underscores the potential of this enzyme (and other diaminopimelate dehydrogenases) as a promising candidate for supporting the enzymatic production of halogenated D-amino acids.

## CONCLUSIONS

The present study provides new insights on the enzymatic production of fluorinated versions of Ala through reductive amination. As the biocatalysis toolbox for the production of alternative building blocks, e.g. NCAAs, continues to expand [58–60], approaches for the selective production of FAA enantiomers (especially, Ala [61]) are increasingly needed. In this work, a kinetic characterization of *^St^*DADPH against FPyr and *^Vp^*ALDH against both FPyr and F3Pyr showed that the monofluorinated substrate decreased the reaction rate with little effect on the enzyme affinity, whereas the trifluorinated version of Pyr exhibited a strong detrimental impact on both affinity and reaction velocity. The production of FAla was combined with the simultaneous regeneration of the reduced cofactor through the oxidation of inexpensive formate by NAD(P)-*^Pse^*FDH. The implementation of this efficient regeneration system led to high conversion yields and enantiomeric purity for both (*S*)-FAla and (*R*)-FAla. *^Sl^*AlaR was determined to be an enzyme with an unexpected high efficiency in dehalogenating FAla through β-elimination, suggesting potential applications in the field of bioremediation [62]. Taken together, our results provide novel approaches to the synthesis of non-canonical building blocks for life [24], which constitutes a decisive step to engineering living cells with alternative lifestyles and functions [63].

## METHODS

### Chemicals and reagents

(*S*)-3-Fluoroalanine and (*R*)-3-fluoroalanine were purchased from BLD Pharmatech Ltd. (Shanghai, China). Trifluoropyruvic acid was acquired from Fluorochem Ltd. (Glossop, UK), isopropyl-β-D-1-thiogalactopyranoside (IPTG) was purchased from Biosynth AG (Staad, Switzerland). All other reagents were acquired from Sigma-Aldrich Co. (St. Louis, MO, USA) unless otherwise specified.

### DNA synthesis, protein production and purification

The list of proteins produced and purified in this study comprise an Ala dehydrogenase from *Vibrio proteolyticus* (*^Vp^*ALDH, UniProt ID O85596), a diaminopimelate dehydrogenase from *Symbiobacterium thermophilum* (*^St^*DAPDH, UniProt ID Q67PI3), an Ala racemase from *Streptomyces lavendulae* (*^Sl^*AlaR, UniProt ID Q65YW7), an Ala racemase from *Escherichia coli* (*^Ec^*AlaR, UniProt ID P0A6B4), a NAD^+^-dependent formate dehydrogenase (FDH) from *Pseudomonas* sp. 101 (i.e. NAD-*^Pse^*FDH) and its site-specific engineered variant (originally named FDH V9) [40], which is characterized by a shifted specificity towards NADP^+^ as a cofactor [64]. For the sake of simplicity, the latter was termed NADP-*^Pse^*FDH throughout this study. The gene fragments encoding *^Vp^*ALDH, *^St^*DAPDH, *^Sl^*AlaR and *^Ec^*AlaR were codon-optimized for expression in *E. coli* K12 (and other Gram-negative bacteria) as explained elsewhere [65] and synthesized *de novo* by Twist Bioscience (San Francisco, CA, USA). Plasmids pZ-ASL [40] carrying either the coding sequence of the parental NAD-*^Pse^*FDH or its engineered version were kindly provided by the Bar-Even group (Max Planck Institute of Molecular Plant Physiology, Golm, Germany). The selected genes were incorporated into a modified pET28a(+) vector (Novagen^TM^, Sigma-Aldrich Co.) encoding the tobacco etch virus (TEV) protease-cleavage site instead of the conventional recognition motif for thrombin protease; the relevant coding sequences were amplified by PCR from the synthetic DNA and the final constructs were obtained through uracil excision (*USER*) cloning [66]. To this end, USER-primers were designed using the AMUSER tool [67]; the native *START* codon of the genes indicated above was removed and the DNA sequence was modified to encode a protein carrying a *N*-terminal His6×-tag followed by the TEV site. The codon-optimized gene sequences and the oligonucleotides needed for USER assembly are listed in **Supplementary Tables S1** and **S2**; the cloning procedures used in this study followed well-established protocols [68–71]. Chemically-competent *E. coli* DH5α λ*pir* cells [72] were used for routinary gene cloning and plasmid construction, and the resulting constructs were sequence-verified prior to be transferred into the expression host *E. coli* BL21(DE3) [F^−^ λ^−^ *ompT hsdSB*(rB^−^, mB^−^) *gal dcm* (DE3); Thermo Fisher Co., San Jose, CA, USA] for protein production.

Protein production experiments started with the preparation of 10-mL precultures by inoculating single colonies of the corresponding recombinant *E. coli* BL21(DE3) strains. Precultures were prepared in 2×YT medium [73] supplemented with 50 μg mL^−1^ kanamycin and incubated overnight at 37°C with agitation at 200 rpm. Then, a 5-mL aliquot of the precultures was used to start 500-mL cultures set in 2-L baffled Erlenmeyer flasks, using 2×YT medium supplemented with 50 μg mL^−1^ kanamycin. These cultures were incubated under the same conditions indicated above until an optical density at 600 nm (OD_600_) of 0.5-0.7 was reached, whereupon the cultures were cooled down to 4°C with no further shaking. Protein expression was induced by adding IPTG to the cultures at a final concentration of 0.4 mM. The induced cultures were incubated at 20°C and 200 rpm for 18 h, then harvested by centrifugation (4,000×*g*, 20 min, 4°C) and stored at –20°C until further processing. Cell lysis was done by sonication, resuspending the frozen bacterial pellets in 10 mL of buffer A (20 mM sodium phosphate pH = 7.5, 300 mM NaCl and 20 mM imidazole) and then submitting the suspension to 2 rounds of ultrasound treatment using a Vibra-Cell^TM^ instrument (model VCX130; Sonics & Materials Co., Newtown, CT, USA) equipped with a 6-mm probe (ref. 630-0422). The procedure consisted of two 7 min-series of sonication at 50% amplitude through 30 s/30 s ON/OFF cycles while keeping the samples on ice to prevent overheating. The lysates were then treated with 2.5 U mL^−1^ Pierce™ universal nuclease for cell lysis (Thermo Fisher Scientific Co.) at room temperature with gentle shaking for 30 min, followed by 20 min centrifugation at 12,000×*g* and 4°C to remove cell debris. The resulting supernatants were filtered through 0.2 μm-membranes and the His6×-tagged enzymes were purified by means of immobilized metal chelate affinity chromatography [74]. Purification was performed using 1 mL of Ni^2+^-nitriloacetic acid (NTA) resin (HisPur^TM^ Ni-NTA resin, Thermo Fisher Scientific Co.) loaded onto 10 mL-Pierce™ disposable columns. The resin was equilibrated in 10 mL of buffer A, then the corresponding clear lysate was passed through the resin, followed by a washing step with 20 mL of buffer A. The bound proteins were eluted by applying 4 mL of buffer B (20 mM sodium phosphate, pH = 7.5, 300 mM NaCl and 500 mM imidazole). The next step consisted of exchanging the buffer to C (20 mM sodium phosphate pH = 7.5, 300 mM NaCl and 1 mM EDTA) by centrifuging the purified enzyme preparations (4,000×*g*, 4°C) using Amicon^TM^ Ultra-15 centrifugal filters of 10 kDa-pore size (Merck-MilliPoreSigma, Burlington, MA, USA). The absorbance at 280 nm (A_280_) of purified enzyme preparations was measured in a NanoDrop™ 2000 spectrophotometer (Thermo Fisher Scientific Co.) and the protein concentration was determined based on the respective theoretical molecular weight and the molar extinction coefficient (ε). The purity of the recombinant enzymes was validated by SDS-PAGE analysis, and the enzyme preparations were stored at 4°C until further use.

### Enzyme assays and assembly of *in vitro* enzyme cascades

Unless otherwise stated, the enzymatic activities were determined by following the oxidation of the NAD(P)H cofactor over time. Samples were prepared in triplicates in a final volume of 200 μL and vigorously mixed with the assay mixture. The absorbance at 340 nm (A_340_) was monitored in 50 s-intervals at 30°C in an *EPOCH2* microplate reader (BioTek Instruments, Winooski, VT, USA). Enzymatic activities were determined by comparison with the corresponding blank experiments, without any catalyst added. The composition of the different reaction mixtures included 50 mM sodium phosphate (pH = 8.0), 200 mM NH_4_Cl and 0.1 g L^−1^ bovine serum albumin. The initial concentration of F*n*Pyr, enzyme and NAD(P)H was adjusted depending on the specific requirements of the assay. NADH and NADPH were added to assays containing *^Vp^*ALDH and *^St^*DAPDH, respectively. Pyridoxal 5’-phosphate was added at 50 μM to assays including an AlaR enzyme. Thus, the kinetics for *^Vp^*ALDH were determined using 0.5 mM NADH and a substrate range of 0-2.5 mM for Pyr; 0-10 mM for FPyr and 0-800 mM for F_3_Pyr. The kinetics for *^St^*DAPDH were analyzed against Pyr (0-100 mM) and FPyr (0-120 mM) using 0.5 mM NADPH in all cases. Kinetic data were fit to the classical Michaelis-Menten kinetic model using SigmaPlot 15.0 (Systat Software Inc., San Jose, CA, USA). Reaction samples to be analyzed by LC-MS or NMR were prepared with an increased concentration of NAD(P)H of 2 mM in order to boost the final amount of fluorinated product(s).

In LC-MS analyses, the enzymatic production of FAla was assayed by using 10 mM FPyr and 0.05 μM *^Vp^*ALDH or 20 μM *^St^*DAPDH, as indicated in the specific experiment. The biosynthesis of F_3_Ala was performed by mixing 100 mM F_3_Pyr with 8 μM *^Vp^*ALDH or 150 μM *^St^*DAPDH. The reactions were incubated at 800 rpm and 30°C for 2 h, before stopping the assay by adding methanol at a final concentration of 50% (v/v). In the case of *^St^*DAPDH and when using F_3_Pyr as the substrate, the reaction time was extended to 20 h. For NMR analyses, the synthesis of FAla was carried out by using 10 mM FPyr and 0.05 μM *^Vp^*ALDH or 20 μM *^St^*DAPDH. The enzymatic production of F_3_Ala was performed with 100 mM F_3_Pyr and 8 μM *^Vp^*ALDH or 150 μM *^St^*DAPDH. In all cases, deuterated water (D_2_O) was added at 10% (v/v). The reaction progression was monitored by recording the spectra over time at 30°C as described in the text.

An enzyme cascade was designed to assay the activity of *^Sl^*AlaR against (*R*)-FAla (**Scheme 3**). *^Vp^*ALDH and *^Sl^*AlaR were combined (each at 0.05 μM) with 1 U mL^−1^ D-amino acid oxidase (DAAO) from porcine kidney (Merck product no. A5222) and 2 U mL^−1^ catalase from bovine liver (Merck-MilliPoreSigma, ref. C1345). NADH was added at a final concentration of 1 mM, and FPyr was used as a substrate either at 0.5 mM (limiting concentration compared to NADH) or 5 mM (excess concentration compared to NADH).

The ability of *^Sl^*AlaR and *^Ec^*AlaR to β-eliminate F atoms was assayed directly against (*S*)-FAla and (*R*)-FAla as the substrates. Two different amounts of enzyme, i.e. 0.02 μM and 0.1 μM, were added to 10 mM of each FAla isomer. Reactions were incubated at 30°C and 800 rpm for 18 h and stopped by the end of the incubation through the prompt addition of formic acid at 5% (v/v). The elimination of F was analyzed by HPLC through the quantification of the released Pyr, stoichiometric with respect to F^−^ formed. The number of catalytic cycles per unit of biocatalyst was calculated using two different amounts of enzymes (i.e. 0.02 μM and 0.1 μM) and comparing the amount of molecules of product measured with the number of molecules of AlaR added to the assay.

### Continuous enzymatic production of FAla *in vitro*

Production of FAla from FPyr using either *^Vp^*ALDH or *^St^*DAPDH was combined with (NAD/NADP)-*^Pse^*FDH and formate as a regeneration system for NAD(P)H (**Scheme 4**). FPyr was added at 20 mM, whereas the initial concentration of formate and NH_4_Cl was set to 200 mM. For the production of (*R*)-FAla, *^Vp^*ALDH (5 μM) was combined with NAD-*^Pse^*FDH (5 μM) and 1 mM NADH. For the production of (*S*)-FAla, *^St^*DAPDH (50 μM) was combined with NADP-*^Pse^*FDH (5 μM) and 1 mM NADPH. The reaction samples were incubated at 800 rpm and 30°C for 12 and 24 h and stopped by freezing the reaction mixtures at –80°C. In all cases, the production and the enantiomeric distribution of FAla on each condition were determined by HPLC. The HPLC analysis required a previous derivatization using the Marfey’s reagent (1-fluoro-2,4-dinitrophenyl-5-L-alanine amide, FDAA), which reacts with primary amines allowing for the separation of amino acid enantiomers by reverse-phase chromatography. The reaction samples were diluted 1:25 prior to derivatization, which was performed by following the manufacturer’s instructions (Thermo Fisher Scientific Co.).

### HPLC analysis

The β-elimination of F through the activity of either *^Sl^*AlaR or *^Ec^*AlaR against FAla was analyzed by HPLC, based on the formation of Pyr. The analysis was performed on a Dionex UltiMate^TM^ 3000 HPLC system equipped with a RefractoMax 521 refractive index detector (IDEX Health & Science LLC, Oak Harbor, WA, USA). Samples were loaded onto an Aminex^TM^ HPX-87X ion exclusion (300×7.8 mm) column (Bio-Rad Laboratories, Hercules, CA, USA) kept at 30°C, and the mobile phase was 5 mM H_2_SO_4_ at 0.6 mL min^−1^ with isocratic elution applied for 18 min [75]. A calibration curve of sodium pyruvate was prepared from 0.2 to 12 mM in each set of measurements. Similarly, the production and enantiomeric distribution of FAla was assessed by HPLC after derivatizing the samples. The analysis was performed on a Dionex UltiMate^TM^ 3000 HPLC system equipped with a Dionex UltiMate^TM^ 3000 Diode Array Detector; absorbance was continuously monitored at 310 nm. The quantification of FAla was based on a calibration curve of the corresponding pure enantiomers in a range of concentrations from 0.08 to 1.25 mM. Samples were loaded onto a Supelco™ Discovery™ HS F5-3 (15 cm×2.1 mm, 3-μm) column (Sigma-Aldrich Co.) kept at 30 °C. The mobile phase was composed of 10 mM ammonium formate (pH = 3) and acetonitrile; an isocratic elution with 5% (v/v) acetonitrile was applied for the first 0.5 min, then the concentration of acetonitrile was increased to 60% (v/v) over a 6.5 min-gradient. The mobile phase was kept at 60% (v/v) acetonitrile for 2.5 min and then the system was re-equilibrated to the initial conditions applying an isocratic gradient of 5% (v/v) acetonitrile for 2.5 min. The flow was set to 0.7 mL min^−1^ for the entire run. Data processing was carried out using the Chromeleon™ Chromatography Data System (CDS) software 7.2.9 (Thermo Fisher Scientific Co.).

### Mass spectrometry analysis

LC-MS(MS) analysis was performed using an UltiMate^TM^ 3000 UHPLC binary system coupled to an Orbitrap Fusion^TM^ mass spectrometer (Thermo Fisher Scientific Co.). Compound separation was achieved using a Waters^TM^ Acquity UPLC BEH Amide (10 cm×2.1 mm, 1.7-μm) column equipped with an Acquity UPLC BEH amide guard column kept at 40°C. The mobile phases consisted of MilliQ^TM^ water and 0.1% (v/v) formic acid (buffer A), and acetonitrile and 0.1% (v/v) formic acid (buffer B) at a flow rate of 0.35 mL min^−1^. The elution was done through an initial step composed by 85% buffer B, held for 0.8 min, followed by a linear gradient to 50% buffer B over 3.2 min and held for 1 min and then linearly increased for 1 min to 30% buffer B before going back to initial conditions (the re-equilibration time was 3 min). The injection volume was set at 1 μL. The MS(MS) measurements were done in positive-heated electrospray ionization (HESI) mode with a voltage of 3,500 V acquiring in full MS/MS spectra (data dependent acquisition-driven MS/MS) with a *m/z* range of 50–500. The MS1 resolution was set at 120,000 and the MS2 resolution was set at 30,000. Precursor ions were fragmented by stepped high-energy collision dissociation (HCD) using collision energies of 20, 40 and 55 eV. The automatic gain control (AGC) target value was set at 4×10^5^ for the full MS and 5×10^4^ for the MS/MS spectral acquisition. Data analysis was performed using the FreeStyle 1.8 software (Thermo Fisher Scientific Co.).

### NMR analysis

^19^F-NMR spectra were measured with a Bruker Avance III^TM^ HD spectrometer (Bruker Corp, Billerica, MA, USA) operating at a magnetic field *B* = 18.8 T (ν19F = 752.83 MHz) and equipped with a 5-mm TCI ^2^H/^19^F-^13^C-^15^N CryoProbe [76]. All samples were measured at 30°C unless otherwise noted. Each sample (400 μL) was added to a 5-mm NMR tube, shaken and quickly transferred to the spectrometer. For each spectrum in the time series, 100 free induction decay signals were recorded using a π/6 excitation pulse and 3 s of interscan delay, yielding a data point for every 5 min; 20-Hz sample rotation was applied during the measurements to increase mixing. Data analysis (including baseline correction and peak integration) was performed using the TopSpin 4.1.3 NMR software (Bruker Corp.). Due to the overlap between the multiplet signals from the FAla and FPyr, the total FAla signal intensity was determined by peak simulation of the most deshielded peak (calibrated to –228.760 ppm) of the *ddd* multiplet and multiplying this value by 8. The resulting FAla signal intensity was then subtracted from the full integral covering both FAla and FPyr multiplets to obtain the FPyr signal intensity. For the F_3_Ala samples, simple peak integration was applied (using equal-sized integral regions) to extract the F_3_Ala and F_3_Pyr signal intensities [77].

### Data and statistical analysis

All the experiments reported were independently repeated at least three times (as indicated in the corresponding figure or table legend), and the mean value of the corresponding parameter ± standard deviation is presented unless indicated otherwise. Data analysis was performed with Prism 8 (GraphPad Software Inc., San Diego, CA, USA) unless differently specified.

## ACKNOWLEDGMENTS

M.N.D acknowledges the support received from the European Union’s *Horizon2020* Research and innovation programme under the Marie Sklodowska-Curie grant agreement No. 713683 (*COFUNDfellowsDTU*) and from the VELUX Foundation under the Villum Experiment program (project No. 40979). The financial support from The Novo Nordisk Foundation (NNF10CC1016517 and NNF18CC0033664) and from the European Union’s *Horizon2020* Research and Innovation Program under grant agreement No. 814418 (*SinFonia*) to P.I.N. is gratefully acknowledged. The responsibility of this article lies with the authors; the NNF and the European Union are not responsible for any use that may be made of the information contained herein.

## SCHEMES AND FIGURES

**Scheme 1.**
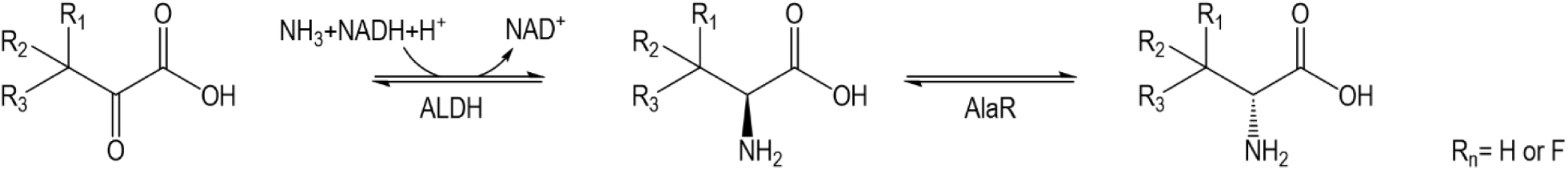
Two-step synthesis of (*S*)-F*_n_*-Ala by the sequential action of ALDH and AlaR.

**Scheme 2.**
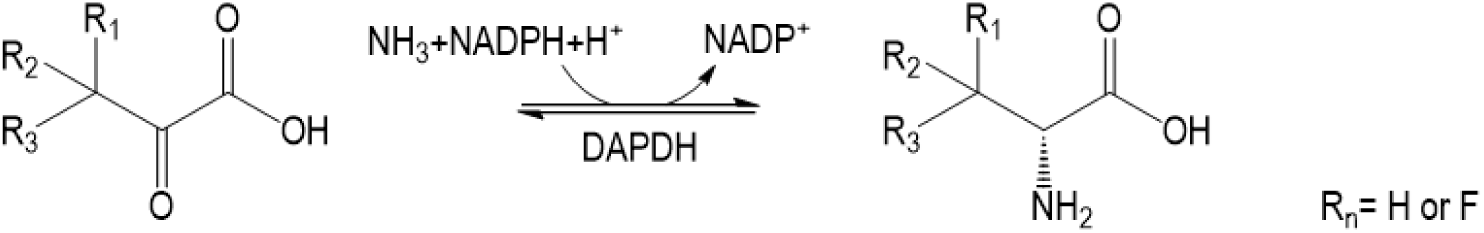
One-step synthesis of (*S*)-F*_n_*-Ala by DAPDH.

**Scheme 3.**
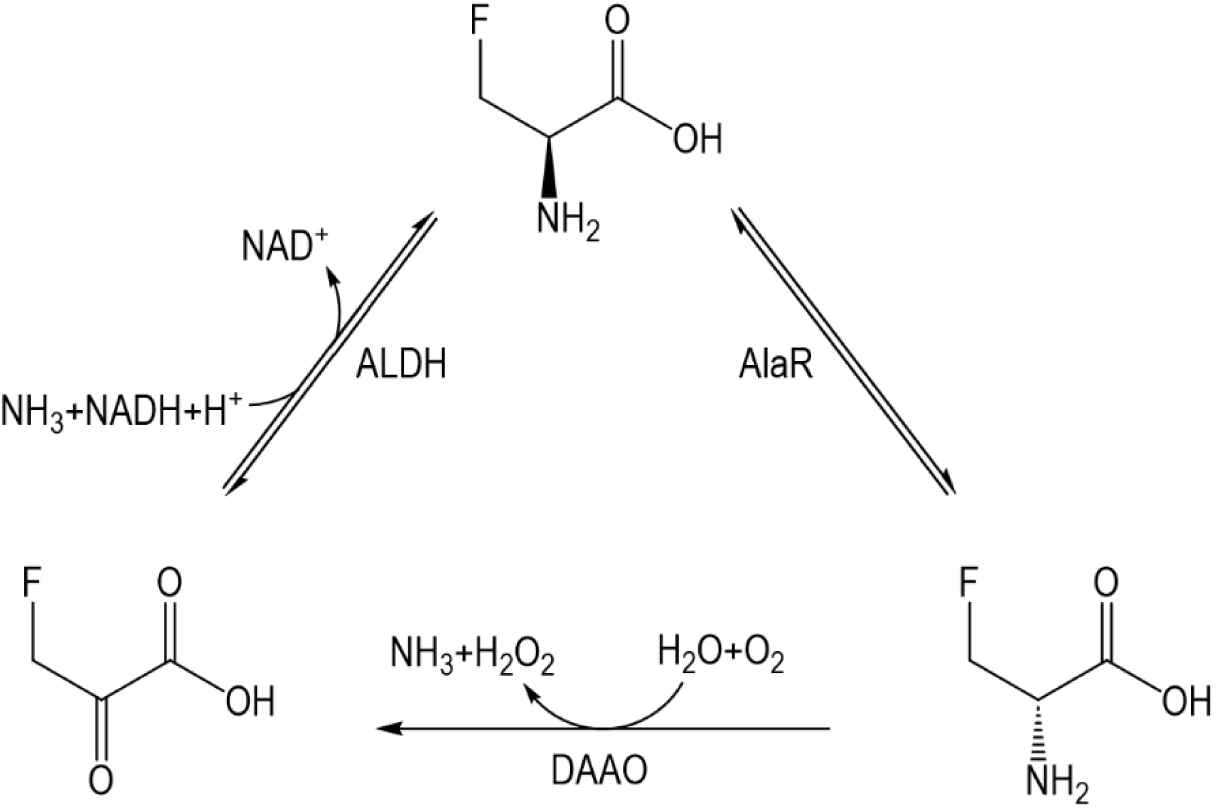
FPyr regeneration by DAAO.

**Scheme 4.**
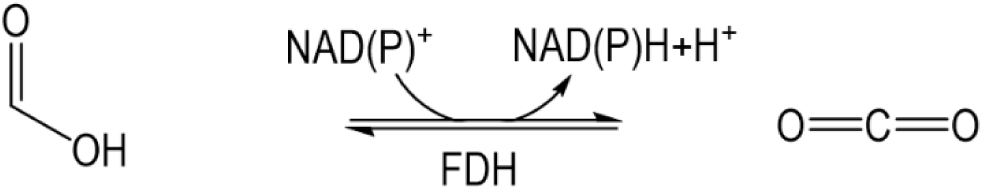
NAD(P)H regeneration by formate dehydrogenase.

## REFERENCES

[1] Clomburg J.M., Crumbley A.M., Gonzalez R. Industrial biomanufacturing: the future of chemical production. Science 2017; 355: aag0804.

[2] Li C.J., Trost B.M. Green chemistry for chemical synthesis. Proc Natl Acad Sci USA 2008; 105: 13197–13202.

[3] Nielsen J., Keasling J.D. Engineering cellular metabolism. Cell 2016; 164: 1185–1197.

[4] Medema M.H., de Rond T., Moore B.S. Mining genomes to illuminate the specialized chemistry of life. Nat Rev Genet 2021; 22: 553–571.

[5] Grigalunas M., Brakmann S., Waldmann H. Chemical evolution of natural product structure. J Am Chem Soc 2022; 144: 3314–3329.

[6] Yang D., Park S.Y., Park Y.S., Eun H., Lee S.Y. Metabolic engineering of *Escherichia coli* for natural product biosynthesis. Trends Biotechnol 2020; 38: 745–765.

[7] Zhou Y., Wang J., Gu Z., Wang S., Zhu W., Aceña J.L., Soloshonok V.A., Izawa K., Liu H. Next generation of fluorine-containing pharmaceuticals, compounds currently in phase II-III clinical trials of major pharmaceutical companies: New structural trends and therapeutic areas. Chem Rev 2016; 116: 422–518.

[8] Wu L., Maglangit F., Deng H. Fluorine biocatalysis. Curr Opin Chem Biol 2020; 55: 119–126.

[9] O’Hagan D. Understanding organofluorine chemistry. An introduction to the C–F bond. Chem Soc Rev 2008; 37: 308–319.

[10] O’Hagan D., Deng H. Enzymatic fluorination and biotechnological developments of the fluorinase. Chem Rev 2015; 115: 634–649.

[11] Buller R., Lutz S., Kazlauskas R.J., Snajdrova R., Moore J.C., Bornscheuer U.T. From nature to industry: harnessing enzymes for biocatalysis. Science 2023; 382: eadh8615.

[12] Walsh C.T., Moore B.S. Enzymatic cascade reactions in biosynthesis. Angew Chem Int Ed Engl 2019; 58: 6846–6879.

[13] Haas R., Nikel P.I. Challenges and opportunities in bringing nonbiological atoms to life with synthetic metabolism. Trends Biotechnol 2023; 41: 27–45.

[14] Cros A., Alfaro-Espinoza G., de Maria A., Wirth N.T., Nikel P.I. Synthetic metabolism for biohalogenation. Curr Opin Biotechnol 2022; 74: 180–193.

[15] Erb T.J., Jones P.R., Bar-Even A. Synthetic metabolism: metabolic engineering meets enzyme design. Curr Opin Chem Biol 2017; 37: 56–62.

[16] Wendisch V.F. Microbial production of amino acids and derived chemicals: synthetic biology approaches to strain development. Curr Opin Biotechnol 2014; 30: 51–58.

[17] Ito T., Muto N., Sakagami H., Tanaka M., Hemmi H., Yoshimura T. D-Amino acid auxotrophic *Escherichia coli* strain for *in vivo* functional cloning of novel D-amino acid synthetic enzyme. FEBS J 2023; 290: 2895–2908.

[18] Gao X., Ma Q., Zhu H. Distribution, industrial applications, and enzymatic synthesis of D-amino acids. Appl Microbiol Biotechnol 2015; 99: 3341–3349.

[19] Pidgeon S.E., Fura J.M., Leon W., Birabaharan M., Vezenov D., Pires M.M. Metabolic profiling of bacteria by unnatural C-terminated D-amino acids. Angew Chem Int Ed Engl 2015; 54: 6158–6162.

[20] Fura J.M., Kearns D., Pires M.M. D-Amino acid probes for penicillin binding protein-based bacterial surface labeling. J Biol Chem 2015; 290: 30540–30550.

[21] Brittain W.D.G., Lloyd C.M., Cobb S.L. Synthesis of complex unnatural fluorine-containing amino acids. J Fluor Chem 2020; 239: 109630.

[22] Nakano S., Kozuka K., Minamino Y., Karasuda H., Hasebe F., Ito S. Ancestral L-amino acid oxidases for deracemization and stereoinversion of amino acids. Commun Chem 2020; 3: 181.

[23] Kawamura Y., Ishida C., Miyata R., Miyata A., Hayashi S., Fujinami D., Ito S., Nakano S. Structural and functional analysis of hyper-thermostable ancestral L-amino acid oxidase that can convert Trp derivatives to D-forms by chemoenzymatic reaction. Commun Chem 2023; 6: 200.

[24] Nieto-Domínguez M., Nikel P.I. Intersecting xenobiology and *neo*-metabolism to bring novel chemistries to life. ChemBioChem 2020; 21: 2551–2571.

[25] Moschner J., Stulberg V., Fernandes R., Huhmann S., Leppkes J., Koksch B. Approaches to obtaining fluorinated α-amino acids. Chem Rev 2019; 119: 10718–10801.

[26] Gonçalves L.P.B., Antunes O.A.C., Pinto G.F., Oestreicher E.G. Kinetic aspects involved in the simultaneous enzymatic synthesis of (*S*)-3-fluoroalanine and (*R*)-3-fluorolactic acid. J Fluorine Chem 2003; 124: 219–227.

[27] Wu L., Deng H. Defluorination of 4-fluorothreonine by threonine deaminase. Org Biomol Chem 2020; 18: 6236–6240.

[28] Humelnicu I., Würthwein E.U., Haufe G. The conformers of 3-fluoroalanine—a theoretical study. Org Biomol Chem 2012; 10: 2084–2093.

[29] Sutherland A., Willis C.L. Synthesis of fluorinated amino acids. Nat Prod Rep 2000; 17: 621–631.

[30] Ohshima T., Wandrey C., Conrad D. Continuous production of 3-fluoro-L-alanine with alanine dehydrogenase. Biotechnol Bioeng 1989; 34: 394–397.

[31] Kato S., Ohshima T., Galkin A., Kulakova L., Yoshimura T., Esaki N. Purification and characterization of alanine dehydrogenase from a marine bacterium, *Vibrio proteolyticus*. J Mol Catal 2003; 23: 373–378.

[32] Hu X., Bai Y., Fan T.P., Zheng X., Cai Y. A novel type alanine dehydrogenase from *Helicobacter aurati*: molecular characterization and application. Int J Biol Macromol 2020; 161: 636–642.

[33] Azam M.A., Jayaram U. Inhibitors of alanine racemase enzyme: a review. J Enzyme Inhib Med Chem 2016; 31: 517–526.

[34] Gao X., Zhang Z., Zhang Y., Li Y., Zhu H., Wang S., Li C. A newly determined member of the meso-diaminopimelate dehydrogenase family with a broad substrate spectrum. Appl Environ Microbiol 2017; 83: e00476–00417.

[35] Liu W., Guo R.T., Chen X., Li Z., Gao X., Huang C.H., Wu Q., Feng J., Zhu D. Structural analysis reveals the substrate-binding mechanism for the expanded substrate specificity of mutant meso-diaminopimelate dehydrogenase. ChemBioChem 2015; 16: 924–929.

[36] Gao X., Huang F., Feng J., Chen X., Zhang H., Wang Z., Wu Q., Zhu D. Engineering the *meso*-diaminopimelate dehydrogenase from *Symbiobacterium thermophilum* by site saturation mutagenesis for D-phenylalanine synthesis. Appl Environ Microbiol 2013; 79: 5078–5081.

[37] Noda M., Kawahara Y., Ichikawa A., Matoba Y., Matsuo H., Lee D.G., Kumagai T., Sugiyama M. Self-protection mechanism in D-cycloserine-producing *Streptomyces lavendulae*. Gene cloning, characterization, and kinetics of its alanine racemase and D-alanyl-D-alanine ligase, which are target enzymes of D-cycloserine. J Biol Chem 2004; 279: 46143–46152.

[38] Badet B., Roise D., Walsh C.T. Inactivation of the *dadB Salmonella typhimurium* alanine racemase by D and L isomers of b-substituted alanines: kinetics, stoichiometry, active site peptide sequencing, and reaction mechanism. Biochemistry 1984; 23: 5188–5194.

[39] Wang E., Walsh C. Suicide substrates for the alanine racemase of *Escherichia coli* B. Biochemistry 1978; 17: 1313–1321.

[40] Calzadiaz-Ramírez L., Calvó-Tusell C., Stoffel G.M.M., Lindner S.N., Osuna S., Erb T.J., Garcia-Borràs M., Bar-Even A., Acevedo-Rocha C.G. *In vivo* selection for formate dehydrogenases with high efficiency and specificity toward NADP^+^. ACS Catal 2020; 10: 7512–7525.

[41] Schröder I., Vadas A., Johnson E., Lim S., Monbouquette H.G. A novel archaeal alanine dehydrogenase homologous to ornithine cyclodeaminase and μ-crystallin. J Bacteriol 2004; 186: 7680–7689.

[42] Zhou M., Feng Z., Zhang X. Recent advances in the synthesis of fluorinated amino acids and peptides. Chem Commun 2023; 59: 1434–1448.

[43] Aceña J.L., Sorochinsky A.E., Soloshonok V.A. Recent advances in the asymmetric synthesis of α-(trifluoromethyl)-containing α-amino acids. Synthesis 2012; 44: 1591–1602.

[44] Ågren D., Stehr M., Berthold C.L., Kapoor S., Oehlmann W., Singh M., Schneider G. Three-dimensional structures of apo– and holo-L-alanine dehydrogenase from *Mycobacterium tuberculosis* reveal conformational changes upon coenzyme binding. J Mol Biol 2008; 377: 1161–1173.

[45] Maeno M., Ohmori T., Nukada D., Sakuraba H., Satomura T., Ohshima T. Two different alanine dehydrogenases from *Geobacillus kaustophilus*: their biochemical characteristics and differential expression in vegetative cells and spores. Biochim Biophys Acta Proteins Proteom 2023; 1871: 140904.

[46] Berkowitz D.B., Karukurichi K.R., de la Salud-Bea R., Nelson D.L., McCune C.D. Use of fluorinated functionality in enzyme inhibitor development: mechanistic and analytical advantages. J Fluor Chem 2008; 129: 731–742.

[47] Chan P.W., Yakunin A.F., Edwards E.A., Pai E.F. Mapping the reaction coordinates of enzymatic defluorination. J Am Chem Soc 2011; 133: 7461–7468.

[48] Thornberry N.A., Bull H.G., Taub D., Wilson K.E., Giménez-Gallego G., Rosegay A., Soderman D.D., Patchett A.A. Mechanism-based inactivation of alanine racemase by 3-halovinylglycines. J Biol Chem 1991; 266: 21657–21665.

[49] Esaki N., Walsh C.T. Biosynthetic alanine racemase of *Salmonella typhimurium*: purification and characterization of the enzyme encoded by the *alr* gene. Biochemistry 1986; 25: 3261–3267.

[50] Roise D., Soda K., Yagi T., Walsh C.T. Inactivation of the Pseudomonas striata broad specificity amino acid racemase by D and L isomers of β-substituted alanines: kinetics, stoichiometry, active site peptide, and mechanistic studies. Biochemistry 1984; 23: 5195–5201.

[51] Deng H., O’Hagan D., Schaffrath C. Fluorometabolite biosynthesis and the fluorinase from *Streptomyces cattleya*. Nat Prod Rep 2004; 21: 773–784.

[52] Weeks A.M., Keddie N.S., Wadoux R.D., O’Hagan D., Chang M.C.Y. Molecular recognition of fluorine impacts substrate selectivity in the fluoroacetyl-CoA thioesterase FlK. Biochemistry 2014; 53: 2053–2063.

[53] McMurry J.L., Chang M.C.Y. Fluorothreonyl-tRNA deacylase prevents mistranslation in the organofluorine producer *Streptomyces cattleya*. Proc Natl Acad Sci USA 2017; 114: 11920–11925.

[54] Kozaeva E., Nieto-Domínguez M., Hernández A.D., Nikel P.I. Synthetic metabolism for *in vitro* acetone biosynthesis driven by ATP regeneration. RSC Chem Biol 2022; 3: 1331–1341.

[55] Mordhorst S., Andexer J.N. Round, round we go—Strategies for enzymatic cofactor regeneration. Nat Prod Rep 2020; 37: 1316–1333.

[56] Yi J., Li Z. Artificial multi-enzyme cascades for natural product synthesis. Curr Opin Biotechnol 2022; 78: 102831.

[57] Serov A.E., Popova A.S., Fedorchuk V.V., Tishkov V.I. Engineering of coenzyme specificity of formate dehydrogenase from *Saccharomyces cerevisiae*. Biochem J 2002; 367: 841–847.

[58] Miller D.C., Athavale S.V., Arnold F.H. Combining chemistry and protein engineering for new-to-nature biocatalysis. Nat Synth 2022; 1: 18–23.

[59] Pyser J.B., Chakrabarty S., Romero E.O., Narayan A.R.H. State-of-the-art biocatalysis. ACS Cent Sci 2021; 7: 1105–1116.

[60] Orton H.W., Qianzhu H., Abdelkader E.H., Habel E.I., Tan Y.J., Frkic R.L., Jackson C.J., Huber T., Otting G. Through-space scalar ^19^F-^19^F couplings between fluorinated noncanonical amino acids for the detection of specific contacts in proteins. J Am Chem Soc 2021; 143: 19587–19598.

[61] Kubyshkin V., Budisa N. Anticipating alien cells with alternative genetic codes: away from the alanine world! Curr Opin Biotechnol 2019; 60: 242–249.

[62] Dvořák P., Nikel P.I., Damborský J., de Lorenzo V. *Bioremediation 3.0*: Engineering pollutant-removing bacteria in the times of systemic biology. Biotechnol Adv 2017; 35: 845–866.

[63] Kubyshkin V., Davis R., Budisa N. Biochemistry of fluoroprolines: the prospect of making fluorine a bioelement. Beilstein J Org Chem 2021; 17: 439–460.

[64] Volke D.C., Martino R.A., Kozaeva E., Smania A.M., Nikel P.I. Modular (de)construction of complex bacterial phenotypes by CRISPR/nCas9-assisted, multiplex cytidine base-editing. Nat Commun 2022; 13: 3026.

[65] Pardo I., Bednar D., Calero P., Volke D.C., Damborský J., Nikel P.I. A nonconventional Archaeal fluorinase identified by *in silico* mining for enhanced fluorine biocatalysis. ACS Catal 2022; 12: 6570–6577.

[66] Cavaleiro A.M., Kim S.H., Seppälä S., Nielsen M.T., Nørholm M.H. Accurate DNA assembly and genome engineering with optimized uracil excision cloning. ACS Synth Biol 2015; 4: 1042–1046.

[67] Genee H.J., Bonde M.T., Bagger F.O., Jespersen J.B., Sommer M.O.A., Wernersson R., Olsen L.R. Software-supported *USER* cloning strategies for site-directed mutagenesis and DNA assembly. ACS Synth Biol 2015; 4: 342–349.

[68] Wirth N.T., Kozaeva E., Nikel P.I. Accelerated genome engineering of *Pseudomonas putida* by I-*Sce*I―mediated recombination and CRISPR-Cas9 counterselection. Microb Biotechnol 2020; 13: 233–249.

[69] Volke D.C., Friis L., Wirth N.T., Turlin J., Nikel P.I. Synthetic control of plasmid replication enables target– and self-curing of vectors and expedites genome engineering of *Pseudomonas putida*. Metab Eng Commun 2020; 10: e00126.

[70] Volke D.C., Olavarría K., Nikel P.I. Cofactor specificity of glucose-6-phosphate dehydrogenase isozymes in *Pseudomonas putida* reveals a general principle underlying glycolytic strategies in bacteria. mSystems 2021; 6: e00014–21.

[71] Fernández-Cabezón L., Cros A., Nikel P.I. Spatiotemporal manipulation of the mismatch repair system of *Pseudomonas putida* accelerates phenotype emergence. ACS Synth Biol 2021; 10: 1214–1226.

[72] Platt R., Drescher C., Park S.K., Phillips G.J. Genetic system for reversible integration of DNA constructs and *lacZ* gene fusions into the *Escherichia coli* chromosome. Plasmid 2000; 43: 12–23.

[73] Sambrook J., Russell D.W., 2001. Molecular cloning: a laboratory manual, 3rd ed. Cold Spring Harbor Laboratory.

[74] Gurdo N., Taylor Parkins S.K., Fricano M., Wulff T., Nielsen L.K., Nikel P.I. Protocol for absolute quantification of proteins in Gram-negative bacteria based on QconCAT-based labeled peptides. STAR Protoc 2023; 4: 102060.

[75] Wirth N.T., Gurdo N., Krink N., Vidal-Verdú A., Donati S., Fernández-Cabezón L., Wulff T., Nikel P.I. A synthetic C2 auxotroph of *Pseudomonas putida* for evolutionary engineering of alternative sugar catabolic routes. Metab Eng 2022; 74: 83–97.

[76] Calero P., Volke D.C., Lowe P.T., Gotfredsen C.H., O’Hagan D., Nikel P.I. A fluoride-responsive genetic circuit enables *in vivo* biofluorination in engineered *Pseudomonas putida*. Nat Commun 2020; 11: 5045.

[77] Smith A.J.R., York R., Uhrín D., Bell N.G.A. New ^19^F NMR methodology reveals structures of molecules in complex mixtures of fluorinated compounds. Chem Sci 2022; 13: 3766–3774.

