## Supplementary Material for "High-yield enzymatic synthesis of mono– and trifluorinated alanine enantiomers"

#### SUPPLEMENTARY METHODS

**Table S1.** Nucleotide sequences of synthetic gene fragments encoding the enzymes of this work.

| Description | Sequence (5'→3') |
|---|---|
| $^{Vp}$*alDH*<br><br>Encoding the alanine dehydrogenase from *Vibrio proteolyticus*<br><br>(UniProt ID O85596) | ATCATCGGTGTTCCGAAAGAAATCAAAAACCACGAATACCGTGTTGGTATGAT<br>CCCGGCGTCTGTTCGTGAACTGATCTCTCACGGTCACCAGGTTTTCGTTGAAA<br>CCAACGCGGGTGCGGGTATCGGTTTCTCTGACGACGACTACATCGCGGTTGGT<br>GCGTCTATCCTGCCGACCGCGGCGGAAGTTTTCGCGCAGGCGGACATGATCGT<br>TAAAGTTAAAGAACCGCAGGCGGTTGAACGTGCGATGCTGAAAGAAGGTCAGA<br>TCCTGTTCACCTACCTGCACCTGGCGCCGGACTTCCCGCAGACCGAAGACCTG<br>ATCAAATCTAAAGCGGTTTGCATCGCGTACGAAACCGTTACCGACAACATGGG<br>TCGTCTGCCGCTGCTGGCGCCGATGTCTGAAGTTGCGGGTCGTATGTCTATCC<br>AGGCGGGTGCGCAGACCCTGGAAAAATCTCACGGTGGTCGTGGTCTGCTGCTG<br>GGTGGTGTTCCGGGTGTTGAACCGGCGAAAGTTGTTATCGTTGGTGGTGGTGT<br>TGTTGGTGCGAACGCGGCGCGTATGGCGGTTGGTATGCGTGCGGACGTTACCA<br>TCCTGGACCGTAACATCGACACCCTGCGTAAACTGGACGAAGAATTCCAGGGT<br>CGTGCGAAAGTTGTTTACTCTACCGAAGACGCGATCGAAAAACACGTTCTGGC<br>GGCGGACCTGGTTATCGGTGCGGTTCTGATCCCGGGTGCGGCGGCGCCGAAAC<br>TGGTTACCAAAGAACACATCGCGAAAATGAAACCGGGTGCGGCGGTTGTTGAC<br>GTTGCGATCGACCAGGGTGGTTGCTTCGAAACCTCTCACGCGACCACCCACGC<br>GGACCCGACCTACATCGTTGACGACGTTGTTCACTACTGCGTTGCGAACATGC<br>CGGGTGCGGTTGCGCGTACCTCTACCTTCGCGCTGAACAACGCGACCCTGCCG<br>TACATCGTTAAACTGGCGAACAAAGGTTACCGTGAAGCGCTGCTGGCGGACCA<br>CGGTTTCCTGGAAGGTCTGAACGTTATCCACGGTAAAGTTACCTGCAAAGAAG<br>TTGCGGAAGCGTTCAACCTGGAATACGTTCAGCCGGAAACCGCGATCGCGATG<br>TTCAACTAA |
| $^{St}$*dapDH*<br><br>Encoding the diaminopimelate dehydrogenase from *Symbiobacterium thermophilum*<br><br>(UniProt ID Q67PI3) | GACAAACTGCGTGTTGCGGTTGTTGGTTACGGTAACGTTGGTCGTTACGCGCT<br>GGAAGCGGTTCAGGCGGCGCCGGACATGGAACTGGTTGGTGTTGTTCGTCGTA<br>AAGTTCTGGCGGCGACCCCGCCGGAACTGACCGGTGTTCGTGTTGTTACCGAC<br>ATCTCTCAGCTGGAAGGTGTTCAGGGTGCGCTGCTGTGCGTTCCGACCCGTTC<br>TGTTCCGGAATACGCGGAAGCGATGCTGCGTCGTGGTATCCACACCGTTGACT<br>CTTACGACATCCACGGTGACCTGGCGGACCTGCGTCGTCGTCTGGACCCGGTT<br>GCGCGTGAACACGGTGCGGCGGCGGTTATCTCTGCGGGTTGGGACCCGGGTAC<br>CGACTCTATCATCCGTGCGCTGCTGGAATTCATGGCGCCGAAAGGTATCACCT<br>ACACCAACTTCGGTCCGGGTATGTCTATGGGTCACTCTGTTGCGGTTAAAGCG<br>ATCCCGGGTGTTCGTGACGCGCTGTCTATGACCATCCCGGCGGGTATGGGTGT<br>TCACAAACGTGCGGTTTACGTTGAACTGGAACCGGGTGCGGACTTCGCGGAAG<br>TTGAACGTGCGATCAAAACCGACCCGTACTTCGTTCGTGACGAAACCCGTGTT<br>ACCCAGGTTGAATCTGTTTCTGCGCTGATGGACGTTGGTCACGGTGTTGTTAT<br>GGAACGTAAAGGTGTTTCTGGTGCGACCCACAACCAGCTGTTCCGTTTCGAAA<br>TGCGTATCAACAACCCGGCGCTGACCGCGCAGGTTATGGTTGCGGCGCTGCGT<br>GCGGCGGCGCGTCAGAAACCGGGTTGCTACACCATGATCGAAATCCCGGTTAT<br>CGACTACCTGCCGGGTGACCGTGAAGCGTGGATCCGTAAACTGGTTTGA |
| $^{Sl}$*alaR*<br><br>Encoding the alanine racemase from *Streptomyces lavendulae*<br><br>(UniProt ID Q65YW7) | AATGAAACACCTACGCGTGTTTACGCCGAGATCGACTTGGACGCCGTTCGTGC<br>TAACGTACGTGCATTGCGCGCCCGTGCACCCCGCTCGGCGCTGATGGCCGTTG<br>TAAAAAGCAATGCTTACGGCCATGGTGCGGTACCATGCGCTCGTGCAGCGCAA<br>GAAGCAGGGGCAGCATGGTTGGGTACGGCCACGCCTGAGGAGGCGTTAGAACT<br>GCGTGCTGCCGGGATCCAGGGTCGCATTATGTGCTGGCTTTGGACCCCGGGTG<br>GGCCATGGCGTGAGGCGATCGAGACTGACATTGATGTAAGTGTGTCAGGTATG<br>TGGGCATTAGATGAAGTTCGTGCAGCAGCGCGCGCAGCTGGCCGTACTGCGCG<br>TATTCAGTTGAAGGCTGATACAGGACTGGGACGTAATGGTTGTCAGCCTGCGG<br>ACTGGGCTGAACTTGTGGGCGCAGCAGTTGCGGCGCAAGCAGAAGGTACGGTG<br>CAGGTTACTGGAGTATGGTCTCATTTCGCTTGCGCTGATGAACCAGGCCACCC<br>ATCGATTCGCTTACAGCTGGATGCGTTCCGCGATATGTTAGCGTACGCAGAAA |

| | |
|---|---|
| | AGGAAGGCGTAGACCCAGAGGTGCGCCATATCGCTAATAGTCCCGCTACTCTG<br>ACCCTTCCAGAGACGCATTTCGATTTAGTACGTACTGGGCTTGCTGTATACGG<br>AGTAAGTCCTTCTCCTGAATTAGGGACGCCGGCGCAATTAGGCCTTCGCCCCG<br>CTATGACACTGCGTGCGAGTCTGGCGTTAGTCAAAACAGTGCCCGCTGGCCAT<br>GGGGTCTCGTATGGTCACCACTATGTCACCGAGTCGGAGACTCACCTGGCTTT<br>AGTACCAGCGGGATACGCAGATGGGATCCCTCGCAACGCCTCCGGTCGCGGGC<br>CTGTCTTAGTGGCGGGTAAGATTCGCCGCGCTGCTGGCCGCATCGCGATGGAT<br>CAATTCGTTGTAGACCTGGGAGAAGATTTGGCAGAGGCTGGCGATGAGGCCGT<br>TATCCTGGGAGATGCAGAACGCGGTGAACCGACTGCTGAGGACTGGGCTCAAG<br>CCGCCCACACAATTGCATATGAAATTGTAACTCGCATCGGTGGCCGCGTTCCA<br>CGCGTTTACCTGGGTGGTTGA |
| $^{Ec}$*alaR*<br><br>Encoding the alanine racemase from *Escherichia coli*<br><br>(UniProt ID P0A6B4) | CAGGCGGCGACCGTTGTTATCAACCGTCGTGCGCTGCGTCACAACCTGCAGCG<br>TCTGCGTGAACTGGCGCCGGCGTCTAAAATGGTTGCGGTTGTTAAAGCGAACG<br>CGTACGGTCACGGTCTGCTGGAAACCGCGCGTACCCTGCCGGACGCGGACGCG<br>TTCGGTGTTGCGCGTCTGGAAGAAGCGCTGCGTCTGCGTGCGGGTGGTATCAC<br>CAAACCGGTTCTGCTGCTGGAAGGTTTCTTCGACGCGCGTGACCTGCCGACCA<br>TCTCTGCGCAGCACTTCCACACCGCGGTTCACAACGAAGAACAGCTGGCGGCG<br>CTGGAAGAAGCGTCTCTGGACGAACCGGTTACCGTTTGGATGAAACTGGACAC<br>CGGTATGCACCGTCTGGGTGTTCGTCCGGAACAGGCGGAAGCGTTCTACCACC<br>GTCTGACCCAGTGCAAAAACGTTCGTCAGCCGGTTAACATCGTTTCTCACTTC<br>GCGCGTGCGGACGAACCGAAATGCGGTGCGACCGAAAAACAGCTGGCGATCTT<br>CAACACCTTCTGCGAAGGTAAACCGGGTCAGCGTTCTATCGCGGCGTCTGGTG<br>GTATCCTGCTGTGGCCGCAGTCTCACTTCGACTGGGTTCGTCCGGGTATCATC<br>CTGTACGGTGTTTCTCCGCTGGAAGACCGTTCTACCGGTGCGGACTTCGGTTG<br>CCAGCCGGTTATGTCTCTGACCTCTTCTCTGATCGCGGTTCGTGAACACAAAG<br>CGGGTGAACCGGTTGGTTACGGTGGTACCTGGGTTTCTGAACGTGACACCCGT<br>CTGGGTGTTGTTGCGATGGGTTACGGTGACGGTTACCCGCGTGCGGCGCCGTC<br>TGGTACCCCGGTTCTGGTTAACGGTCGTGAAGTTCCGATCGTTGGTCGTGTTG<br>CGATGGACATGATCTGCGTTGACCTGGGTCCGCAGGCGCAGGACAAAGCGGGT<br>GACCCGGTTATCCTGTGGGGTGAAGGTCTGCCGGTTGAACGTATCGCGGAAAT<br>GACCAAAGTTTCTGCGTACGAACTGATCACCCGTCTGACCTCTCGTGTTGCGA<br>TGAAATACGTTGACTGA |

**Table S2.** Nucleotide sequences of primers used for plasmid construction.

| Name | Sequence (5'→3')[a] | Description |
|---|---|---|
| pET28a-F | ATCTCTTCuGAGCACCACCACCACC | Amplification of pET-28a(+)-TEV for USER cloning, forward |
| pET28a-R | ATGGCCCuGAAAATAAAGATTCTCGCCGCT | Amplification of pET-28a(+)-TEV for USER cloning with Nt His-tag, reverse |
| $^{vp}$*aldh*-F | AGGGCCAuATCATCGGTGTTCCGAAAGA | Amplification of $^{vp}$*aldh* for USER cloning into pET28a(+)-TEV in-phase with Nt His-tag, removes ATG, forward |
| $^{vp}$*aldh*-R | AGAAGAGAuTTAGTTGAACATCGCGATCG | Amplification of $^{vp}$*aldh* for USER cloning into pET28a(+)-TEV in-phase with Nt His-tag, reverse |
| $^{st}$*dapdh*-F | AGGGCCAuGACAAACTGCGTGTTGCGGT | Amplification of $^{st}$*dapdh* for USER cloning into pET28a(+)-TEV in-phase with Nt His-tag, removes ATG, forward |
| $^{st}$*dapdh*-R | AGAAGAGAuTCAAACCAGTTTACGGATCCA CG | Amplification of $^{st}$*dapdh* for USER cloning into pET28a(+)-TEV in-phase with Nt His-tag, reverse |
| $^{sl}$*alar*-F | AGGGCCAuAATGAAACACCTACGCGTGT | Amplification of $^{sl}$*alar* for USER cloning into pET28a(+)-TEV in-phase with Nt His-tag, removes ATG, forward |
| $^{sl}$*alar*-R | AGAAGAGAuTCAACCACCCAGGTAAACG | Amplification of $^{sl}$*alar* for USER cloning into pET28a(+)-TEV in-phase with Nt His-tag, reverse |
| $^{ec}$*alar*-F | AGGGCCAuCAGGCGGCGACCGTTGTTAT | Amplification of $^{ec}$*alar* for USER cloning into pET28a(+)-TEV in-phase with Nt His-tag, removes ATG, forward |
| $^{ec}$*alar*-R | AGAAGAGAuTCAGTCAACGTATTTCATCGC AACAC | Amplification of $^{ec}$*alar* for USER cloning into pET28a(+)-TEV in-phase with Nt His-tag, reverse |
| $^{pse}$*fdh*-F | AGGGCCAuGCTAAAGTTCTGTGCGTTCTG | Amplification of $^{pse}$*fdh* (both parental and engineered) for USER cloning into pET28a(+)-TEV in-phase with Nt His-tag, removes ATG, forward |
| $^{pse}$*fdh*-R | AGAAGAGAuTTAAACAGCTTTTTTGAATTT AGCAGCTTC | Amplification of $^{pse}$*fdh* (both parental and engineered) for USER cloning into pET28a(+)-TEV in-phase with Nt His-tag, reverse |

[a] The uracil residues relevant for USER cloning are indicated in lowercase.

**Enzyme assays**

The purified racemases were assayed initially against its natural substrate L-Ala (5 mM) to confirm activity. $^{Sl}$AlaR and $^{Ec}$AlaR were added at $4.8\times10^{-3}$ $\mu$M and $1.1\times10^{-2}$ $\mu$M, respectively. The racemase activity was coupled with 1 U $mL^{-1}$ D-amino acid oxidase from porcine kidney (catalogue # A5222, Merck) and 1 U $mL^{-1}$ L-lactic dehydrogenase (catalogue # L2500, Merk). Activity was followed by monitoring the oxidation of 0.5 mM NADH during the conversion of Pyr into L-lactate (**Scheme S1**). Catalase (2 U $mL^{-1}$) from bovine liver (catalogue # C1345, Merck) was added to prevent the accumulation of hydrogen peroxide (**Fig. S2**).

The activity of $^{Pse}$NAD-FDH and $^{Pse}$NADP-FDH was verified against formate. The assay was prepared using 10 mM formate and 50 mM of sodium phosphate (pH = 8.0). For $^{Pse}$NAD-FDH, 0.5 mM $NAD^+$ and $1.8\times10^{-2}$ $\mu$M enzyme were used, whereas the assay for $^{Pse}$NADP-FDH was prepared with 0.5 mM $NADP^+$ and $2.8\times10^{-2}$ $\mu$M biocatalyst. The reduction of the cofactor over time was monitored by the change on absorbance at 340 nm (**Fig. S3**).

#### Supplementary Schemes

**Scheme S1. Racemase activity coupled to lactate synthesis.**

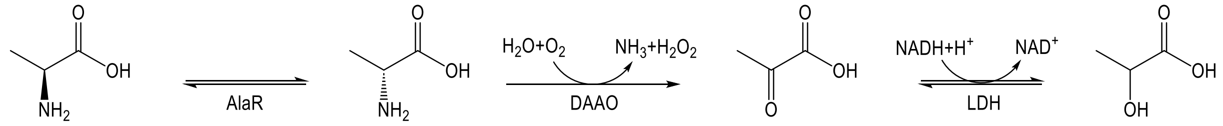

### SUPPLEMENTARY RESULTS

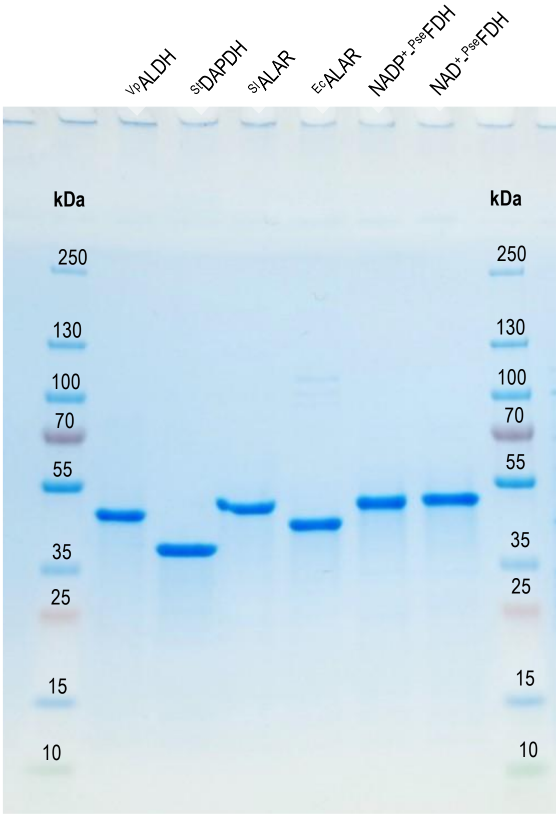


**Fig. S1**. SDS-PAGE of the enzymes produced and purified in this study. A total amount of 1 μg of protein was loaded per lane.

```
VpALDH      MIIGVPKEIKNHEYRVGMIPASVRELISHGHQVFVETNAGAGIGFSDDDYIAVGASILPT 60
MtALDH      MRVGIPTETKNNEFRVAITPAGVAELTRRGHEVLIQAGAGEGSAITDADFKAAGAQLVGT 60
GkALDH      MIIGVPKEIKNNENRVAITPAGVLSFVQAGHTVLIEKEAGVGSGFNDSDYARAGAQIIER 60
            * :*:*.* **:* **.: **.* .:   ** *:::  ** * .:.* *:  .**.::
VpALDH      AAEVFAQADMIVKVKEPQAVERAMLKEGQILFTYLHLAPDFPQTEDLIKSKAVCIAYETV 120
MtALDH      ADQVWADADLLLKVKEPIAAEYGRLRHGQILFTFLHLAASRACTDALLDSGTTSIAYETV 120
GkALDH      AEDVWAQADMVMKVKEPLPSEYRFFRPGLVLFTYLHLAADPELTRVLKESGVIAIAYETV 120
            * :*:*:**:::*****   *   :: * :***:**** .   *  * .* . .******
```

**Fig. S2**. Partial alignment of $^{Vp}$ALDH, the ALDH from *Mycobacterium tuberculosis* ($^{Mt}$ALDH, PDB ID 2VHW_1) and one ALDH from *Geobacillus kaustophilus* ($^{Gk3448}$ALDH, PDB ID 8HYH_1) showing that the residues involved in the pyruvate binding to active site (Arg15, Lys75 and His96) are conserved.

A)

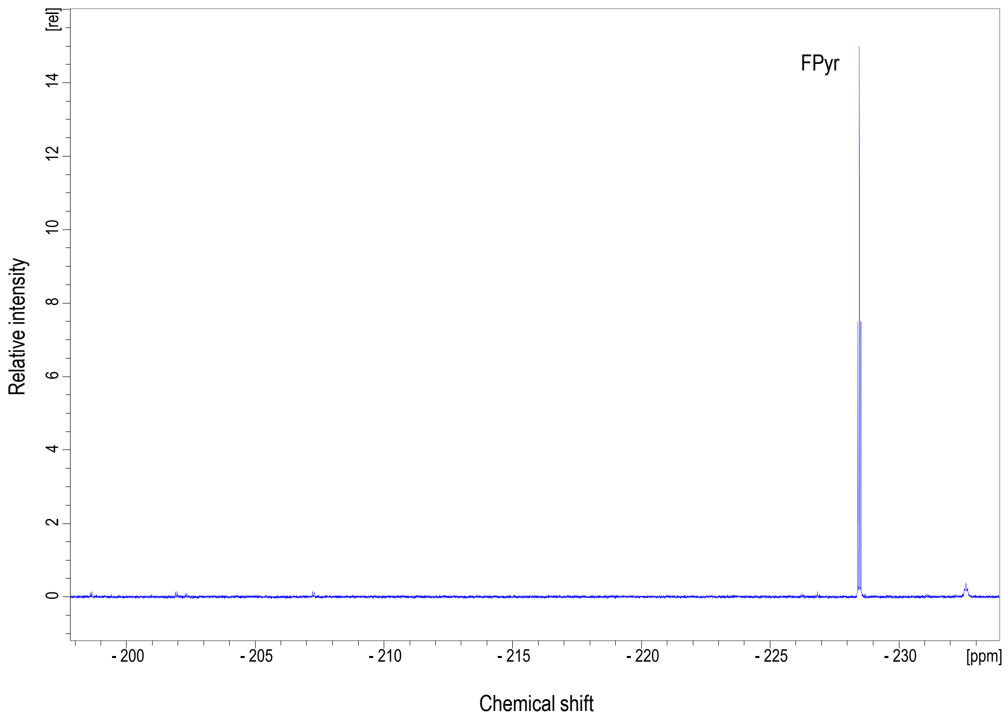


B)

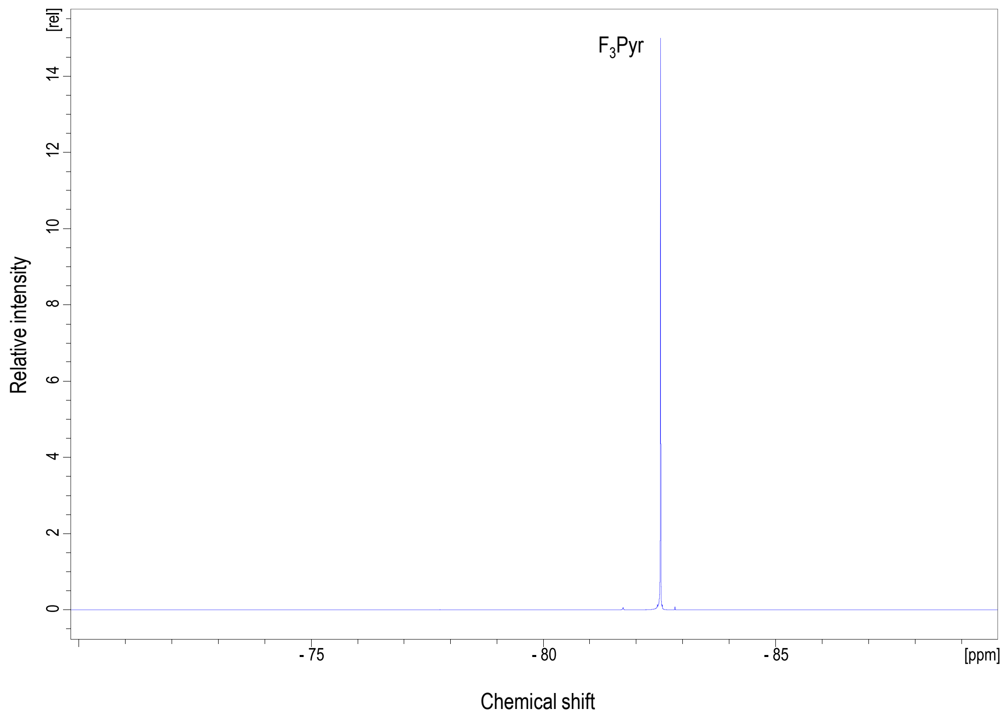


**Fig. S3.** $^{19}$F-NMR spectrum from a reaction mixture containing either (A) FPyr, prior to the addition of $^{Vp}$ALDH or $^{St}$DAPDH to catalyze the corresponding reductive amination (the triplet signal from FPyr is labeled), or (B) $F_3$Pyr prior to the addition of $^{Vp}$ALDH or $^{St}$DAPDH to catalyze the corresponding reductive amination (the singlet signal from $F_3$Pyr is labeled).

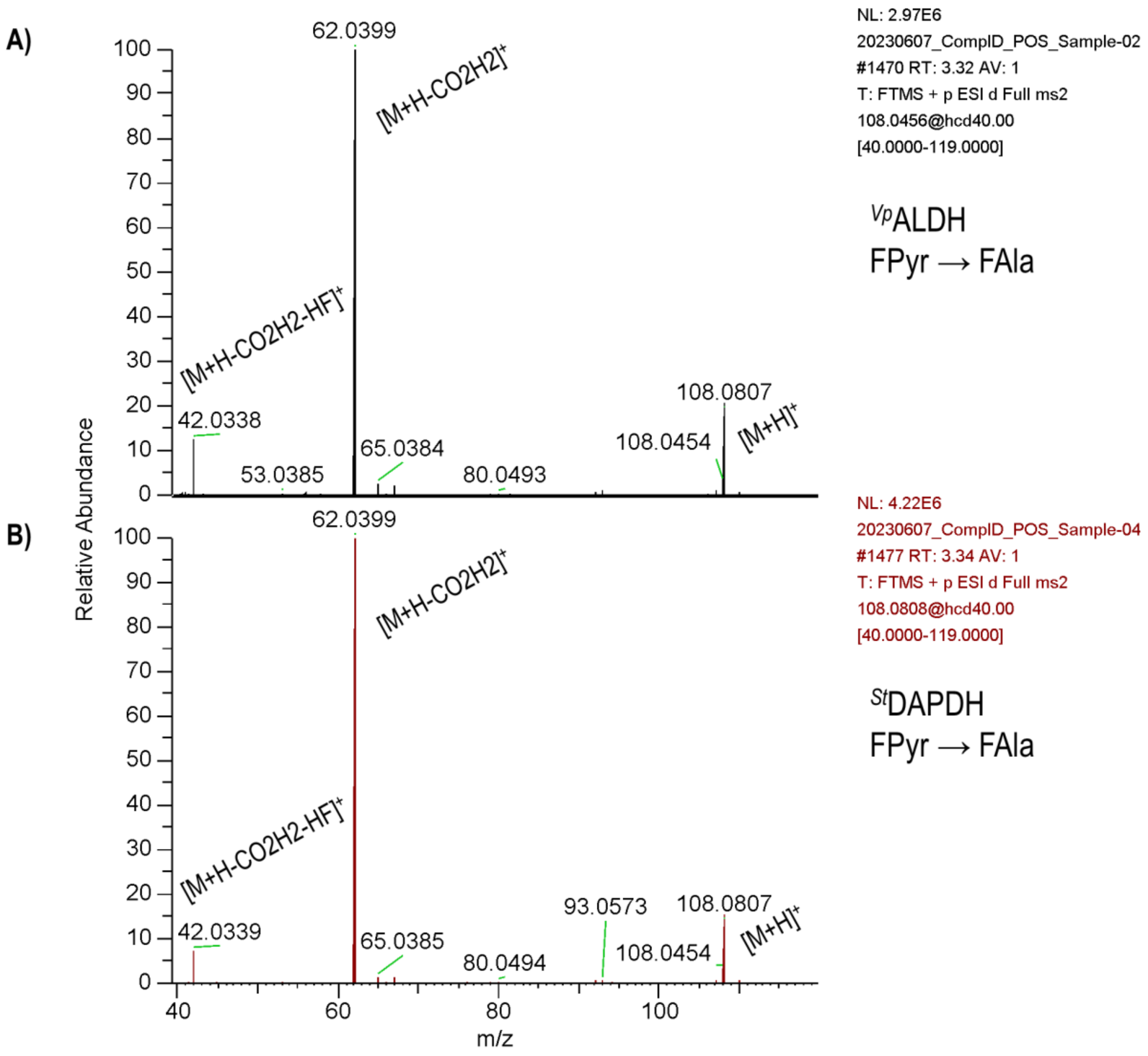


**Fig. S4.** MS(MS) spectra of FAla produced by ${}^{Vp}$ALDH (A) and ${}^{St}$DAPDH (B) from FPyr. Both spectra show the parent ion (*m/z* = 108.08) and the fragments resulting from losing the carboxylic group (*m/z* = 62.04) and both the carboxylic group and the fluorine substituent (*m/z* = 42.03).

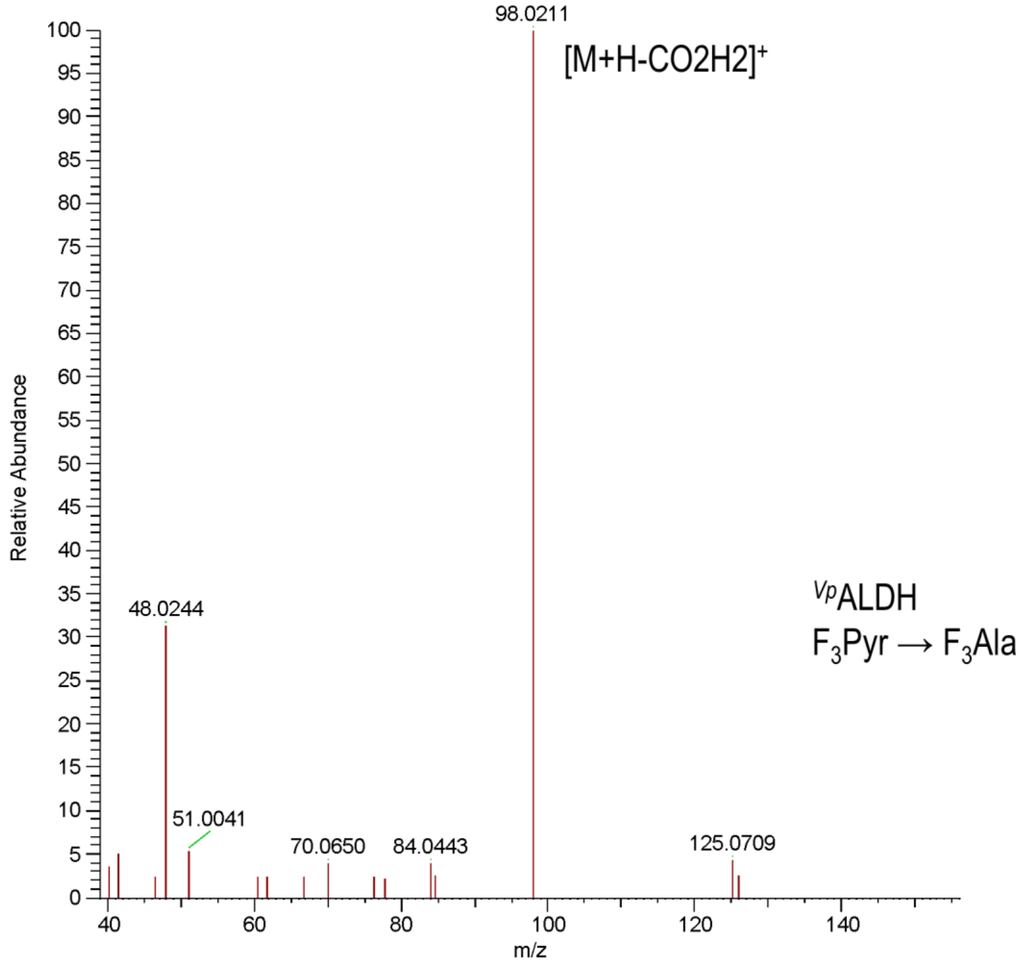


**Fig. S5.** MS(MS) spectrum of $F_3$Ala produced by $^{Vp}$ALDH from $F_3$Pyr. The spectrum displays the daughter ion derived from the loss of the carboxylic group.

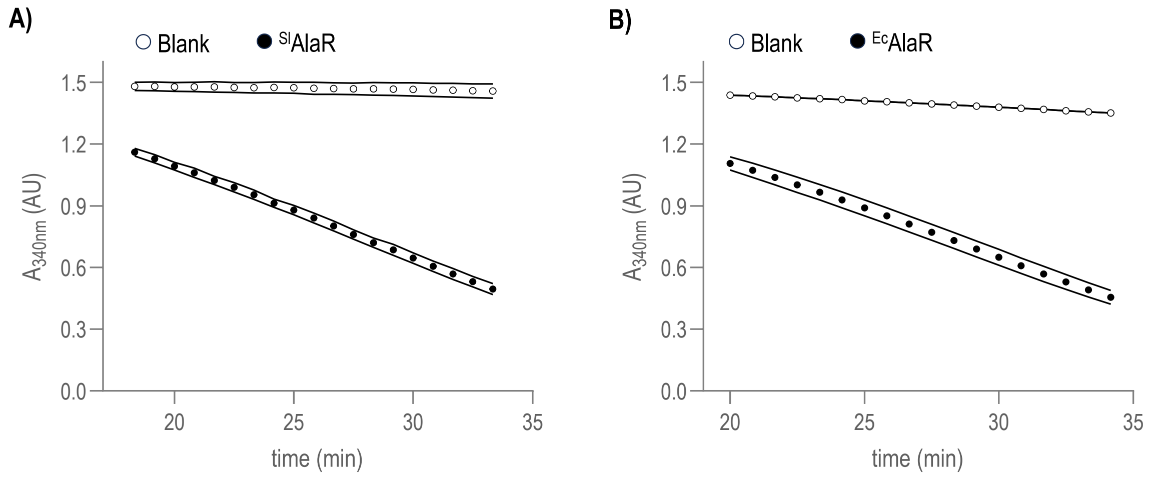


**Fig. S6**. Activity of $^{Sl}$AlaR (A) and $^{Ec}$AlaR (B) against L-Ala over time. The reaction was monitored by coupling the racemase activity with a DAAO and a lactate dehydrogenase. Read-outs show the oxidation of NADH at $A_{340\ nm}$ for both the reaction (*black dots*) and a blank without enzyme (*white dots*). The points represent mean values ± standard deviations from three independent experiments.

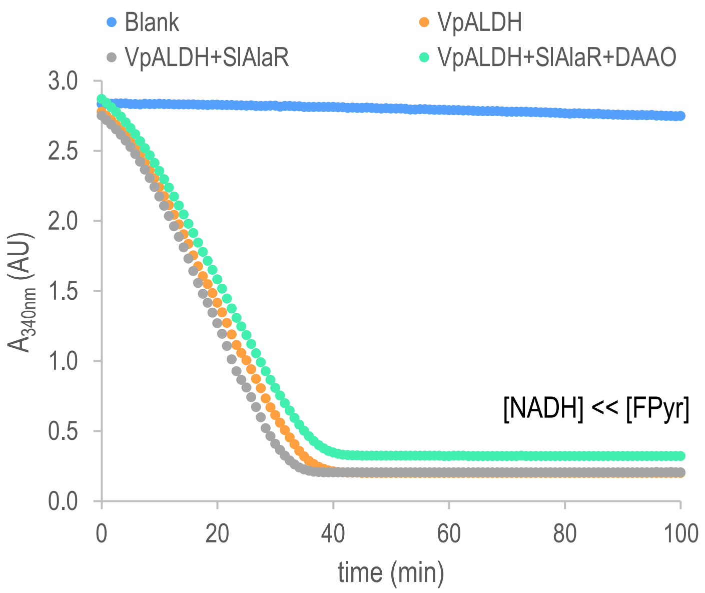


**Fig. S7**. Spectrophotometric monitoring ($A_{340\ nm}$) of NADH oxidation using FPyr in excess compared to the reduced cofactor and different enzyme combinations. The assay serves as a control showing that when FPyr is in excess all the tested combinations reach the reaction completion. The points represent mean values from three independent experiments.

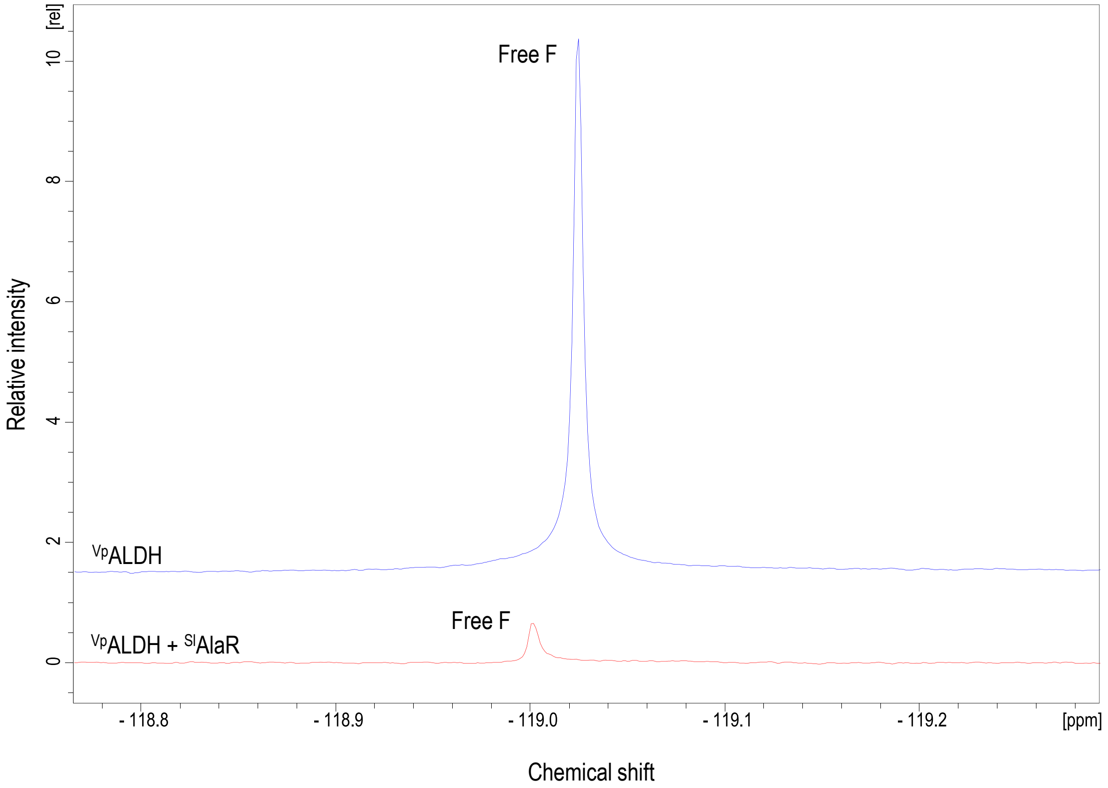


**Fig. S8.** $^{19}F$ NMR spectra showing the relative abundance of free F in equivalent reactions catalyzing the reductive amination of FPyr. The blue spectrum corresponds to an assay containing $^{Vp}$ALDH whereas the red spectrum was obtained from the combination of $^{Vp}$ALDH and $^{Sl}$AlaR. Both were recorded after 100 min of reaction and with the same number of scans.

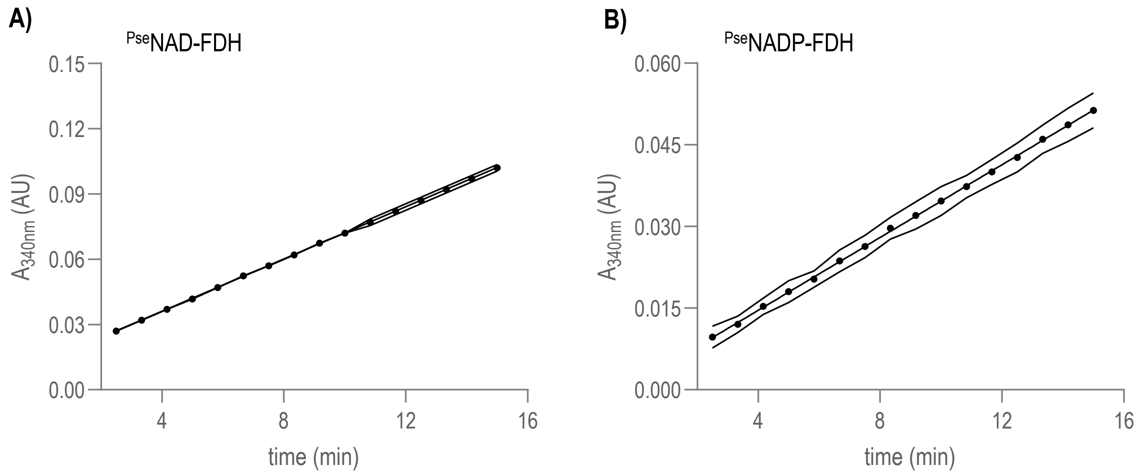


**Fig. S9**. Activity of *Pse*NAD-FDH (A) and *Pse*NADP-FDH (B) against formate over time. Read-outs show reduction of respectively $NAD^+$ and $NADP^+$ at $A_{340}$. The points represent mean values ± standard deviations from three independent experiments.

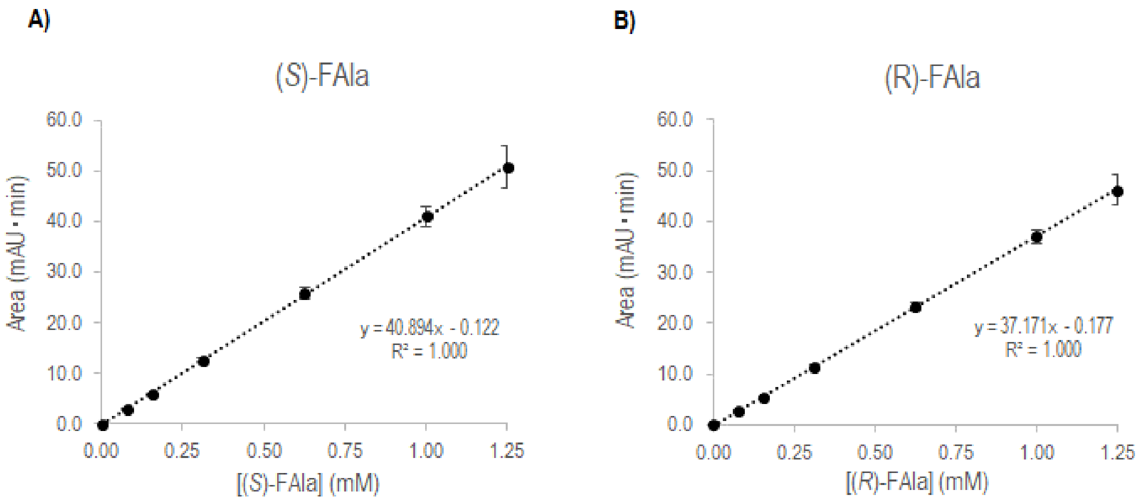


**Fig. S10**. Calibration curves to determine the concentration of the D-Ala (A) and L-Ala (B) enantiomers in the reaction samples. Two series of independent replicates were prepared and analyzed by reversed-phase chromatography. The concentrations of the standards are plotted against the area under the curve. Absorbance was monitored at 310 nm.
